# Cross Potential Selection for Multiple Traits Considering the Progeny Distribution of Future Inbred Lines in Plant Breeding Programs

**DOI:** 10.64898/2026.06.02.729654

**Authors:** Kengo Sakurai, Laurence Moreau, Tristan Mary-Huard, Alain Charcosset, Hiroyoshi Iwata

**Affiliations:** Graduate School of Agricultural and Life Sciences, The University of Tokyo, Bunkyo, Tokyo 113-8657, Japan; Université Paris-Saclay, INRAE, CNRS, AgroParisTech, Génétique Quantitative et Evolution (GQE) - Le Moulon, 91190, Gif-Sur-Yvette, France; Université Paris-Saclay, AgroParisTech, INRAE, UMR MIA Paris-Saclay, 91120, Palaiseau, France

**Keywords:** cross potential selection, multi-trait constraint, progeny distribution, genetic covariance, breeding decision

## Abstract

In plant breeding, it is often necessary to improve a target trait while maintaining other essential traits within desirable ranges. When genetic relationships exist among these traits, improvements in the target trait may lead to undesirable changes in essential traits, complicating cross selections. In such cases, it is critical to select cross-pairs that are expected to produce progeny that satisfy the requirements for all traits. The progeny distribution of each crossing pair can be predicted using the estimated genotypic values and genetic (co)variances of the target and essential traits. By utilizing this distribution, the probability of generating progeny that satisfy predefined trait requirements can be evaluated, allowing a direct comparison of alternative crosses. In this study, we developed Cross Potential Selection for Multiple Traits (CPS-MT), a breeding strategy designed to improve a target trait while maintaining one or more essential traits within desirable ranges. CPS-MT extends the original Cross Potential Selection (CPS) framework to explicitly handle trade-offs between traits under genetic correlations. We evaluated the performance of CPS-MT through simulations involving four types of genetic relationships and two genetic causal factors between traits, resulting in seven scenarios. Across all scenarios, CPS-MT consistently improved the likelihood of obtaining desirable progeny, indicating that CPS-MT provides a practical and effective framework for cross selection under multi-trait constraints in breeding programs.

**Article Summary:** This study developed Cross Potential Selection for Multiple Traits (CPS-MT), a new breeding strategy designed to improve a target trait while maintaining one or more essential traits within desirable ranges. CPS-MT evaluates crossing pairs by predicting progeny distributions based on estimated genotypic values and genetic covariances, enabling direct comparison of alternative crosses under multi-trait constraints. Through simulations incorporating four types of genetic relationships and two causal factors (seven scenarios), CPS-MT consistently increased the likelihood of obtaining progeny that satisfied the predefined trait requirement. These results indicate that CPS-MT provides a practical, robust framework for target trait improvement under trait constraints.

## Introduction

In plant breeding, the genetic improvement of a target trait is achieved through repeated cycles of selection and crossing of promising genotypes. In recent years, genomic prediction (GP) models, which estimate genotypic values using genome-wide marker polymorphisms and phenotypic data (Meuwissen et al. 2001), have attracted considerable attention. GP enables the prediction of genomic estimated breeding values (GEBVs) for untested genotypes based solely on marker information, thereby reducing the need for extensive field evaluation and accelerating breeding cycles.

For self-pollinated crops, an evaluation index for crosses known as the usefulness criterion (UC) has been proposed to quantify the expected value of superior individuals derived from a given cross (Schnell and Utz 1975). UC represents the expected value of superior individuals obtained from a progeny population of a given cross and is defined as UC = *μ* + *ihσ*, where *μ* is the mean GEBV of the cross, *i* is the selection intensity, *h* is the square root of the heritability, and *σ* is the square root of the genetic variance among progeny. Traditionally, *σ* has been calculated by simulating progeny populations and estimating genetic variance from their GEBVs (Mohammadi et al. 2015; Tiede et al. 2015). More recently, theoretical formulas have been developed to calculate *σ* directly using genome-wide marker information, genetic map distances, and estimated marker effects (Lehermeier et al. 2017), with extensions to three-way and four-way crosses (Allier et al. 2019a), and further mathematical improvements enabling rapid computation for large numbers of potential crosses (des Déserts et al. 2023). These advances have made it feasible to systematically evaluate large sets of candidate crosses and have facilitated the development of UC-based breeding strategies. For example, Cross Potential Selection (CPS), which combines UC and a recurrent genomic selection scheme, targets short-term genetic gain (Sakurai et al. 2024), whereas Usefulness Criterion Parental Contribution (UCPC) combines UC and progeny genetic diversity to promote long-term genetic gain (Allier et al. 2019a; Allier et al. 2019b).

The effectiveness of UC-based strategies critically depends on the accuracy of *σ* estimation (Yao et al. 2018), which varies considerably among traits (Neyhart and Smith 2019; Wartha and Lorenz 2024). Accurate estimation of marker effects is therefore crucial for improving the accuracy of *σ* estimation (Lehermeier et al. 2017; Oget-ebrad et al. 2024), and improvement in prediction accuracy can be achieved by increasing the training population size and heritability (Wimmer et al. 2013). For example, in soybean breeding populations, the prediction accuracy of *σ* for yield was reported to be 0.43 in terms of the correlation coefficient (Wartha and Lorenz 2024).

In plant breeding, it is often necessary to improve a target trait while maintaining the genotypic values of other essential traits, that is, traits whose values must remain within predefined desirable ranges. For example, breeders often aim to improve the yield while maintaining the flowering period within a narrow range. When genetic correlations exist between the target and essential traits, selection aimed at improving the target trait may induce undesirable changes in the essential traits (Misztal and Lourenco 2024). Furthermore, genetic relationships between traits are not always linear. For example, yield and plant height are generally positively correlated (Ibrahim 2019; Li et al. 2019; Zhou et al. 2022), whereas excessive increases in plant height often lead to higher lodging scores (Navabi et al. 2006; Wu et al. 2022; Chen et al. 2024). Nonlinear relationships between plant height and yield have been reported for sorghum (Tripathi et al. 2004; Shah et al. 2017), rice (Zhu et al. 2016), and wheat (Addisu et al. 2010; Casebow et al. 2016). Similarly, a nonlinear relationship between days to flowering and yield has been observed in sorghum cultivated in northern Nigeria. Specifically, early flowering lines exhibit reduced light interception, which is essential for photosynthesis, leading to yield reduction, whereas late-flowering lines are more strongly affected by drought stress during the dry season, leading to yield reduction (Flower 1996; Craufurd and Wheeler 2009). These examples indicate that, under certain environmental conditions, essential traits often have an optimal range rather than a monotonic relationship with yield.

Genetic relationships among traits are mainly explained by pleiotropy and spurious pleiotropy. Pleiotropy refers to the phenomenon in which a single gene directly affects multiple traits (Dwivedi et al. 2024). In contrast, spurious pleiotropy arises when two different genes affecting separate traits are located in close proximity and are inherited together because of strong linkage disequilibrium (LD), resulting in an apparent genetic relationship between the traits (Reinert 2022). These distinct causal mechanisms can generate similar phenotypic correlations, leading to different progeny distributions that may substantially influence cross evaluation and selection decisions.

Improving a target trait while controlling changes in other essential traits is a central challenge in plant breeding. When genetic correlations exist between traits, selection aimed solely at improving the target trait may induce unfavorable shifts in essential traits. Despite this challenge, only a limited number of breeding strategies based on genomic information have been developed to address cross selection under multi-trait constraints. One classical approach to this challenge is to use a selection index that combines the GEBVs of multiple traits into a single score using predefined economic weights (Hazel and Lush 1942; Hazel 1943). Multi-Trait Look-Ahead Selection (MT-LAS) is one such strategy designed to improve target traits while maintaining essential traits within desirable ranges (Moeinizade et al. 2020). However, MT-LAS does not account for the progressive fixation of genotypes through repeated selfings, which is a fundamental process in breeding programs for self-pollinated crops, limiting its direct applicability to such crops. The segregation of progeny through repeated selfings is particularly important in self-pollinated crops, as reflected in the long-standing use of UC-based strategies. Therefore, there is a need for a breeding strategy that is applicable to self-pollinated crops and incorporates UC while enabling efficient improvement of a target trait under explicit multi-trait constraints.

In this study, we developed a new breeding strategy that enables the efficient improvement of a target trait while maintaining the genotypic values of one or more essential traits within predefined desirable ranges. Focusing on breeding programs for self-pollinated crops, we designed a recurrent selection framework that promotes genome shuffling and genetic gain (Gaynor et al. 2017) and proposed Cross Potential Selection for Multiple Traits (CPS-MT). CPS-MT extends the single-trait CPS method developed by Sakurai et al. (2024) by explicitly evaluating multivariate progeny distribution, and is formulated to handle one target trait and any number of essential traits. For illustration and detailed performance evaluation, we evaluated CPS-MT in a two-trait setting, where the objective of the breeding program was to improve the genotypic values of Trait1 (target trait) while maintaining those for Trait2 (essential trait) within a predefined desirable range (*l* ≤ *u*_2_ ≤ *h*). To evaluate the effectiveness of CPS-MT, we considered four types of genetic relationships between the two traits (no correlation, positive correlation, and two forms of nonlinear relationships) and assumed two causal mechanisms: pleiotropy and spurious pleiotropy. Seven scenarios were established. For each scenario, 30 independent simulation runs were conducted, and the genotypes were evaluated based on their true genotypic values. As a comparative strategy, CPS-MT, which accounts for progeny genetic (co)variance in cross selection, was compared with Penalized Cross Selection (PCS), which is a baseline approach that evaluates crossing pairs using a weighted combination of mean GEBVs across traits, an extension of the selection index concept to cross evaluation, without incorporating the distribution of progeny genotypic values.

## Materials and methods

### Population and genome data

This section describes the initial population structure and genomic data used in breeding simulations. In this breeding simulation, we randomly selected 150 accessions from a diverse panel of 198 soybean accessions and used them as the initial population (Table S1) (Kajiya-Kanegae et al. 2021). The whole-genome sequencing data encompassed 4,776,813 single-nucleotide polymorphisms (SNPs) distributed across 20 homologous chromosome pairs. SNPs that were heterozygous or had more than 95% missing data, as well as those with a minor allele frequency below 0.01, were excluded. SNPs were filtered based on linkage disequilibrium, retaining SNP pairs with LD < 0.95, resulting in a final set of 303,002 SNP markers. We assumed that each chromosome had a length of one Morgan and that there was a linear relationship between map distances and physical distances. Consequently, linkage map positions were calculated based on the physical positions of adjacent SNPs.

### Marker effects

We set a target trait (Trait1) and an essential trait (Trait2) for the plant breeding programs. We assumed four types of relationships between Trait1 and Trait2: no relationship (“No Relation”), positive relationship (“Positive”), nonlinear relationship 1 (“Nonlinear1”), and nonlinear relationship 2 (“Nonlinear2”). Genetic correlation (*r*_*g*_) between Trait1 and Trait2 was set as *r*_*g*_ = 0 and *r*_*g*_ = 0.6 in “No Relation” and “Positive” cases. In cases of nonlinear relationships, two patterns were assumed regarding Trait2, which exhibits a quadratic relationship with Trait1: one in which the trait value at the optimum of the quadratic relationship lies within the desirable range of Trait2 (“Nonlinear1”), and another in which it lies outside that desirable range (“Nonlinear2”) (Figure S1). A detailed explanation of these nonlinear relationships is provided in Supplementary File S1. Although negative correlations between traits can occur, positive and negative correlations are considered to represent the same relationship, with only a difference in direction. Therefore, negative correlations were excluded in the simulation settings.

Two causes of the genetic relationships between traits were assumed: pleiotropy and spurious pleiotropy. Genetic relationships may arise due to a combination of these factors. However, to facilitate the evaluation of the effectiveness of the proposed method, this study assumed that each genetic relationship arose from one of the two factors. For the uncorrelated case, the underlying causes of trait relationships were not considered. As a result, a total of seven scenarios were assumed: three types of genetic relationships (“Positive,” “Nonlinear1,” and “Nonlinear2”) × two underlying causes (pleiotropy and spurious pleiotropy), plus one uncorrelated case (Figure 1). True marker effects for Trait1 (***β***_(1)_) and Trait2 (***β***_(2)_) were generated according to the scenario settings in each independent breeding simulation. A detailed explanation of the assignment of marker effects and the genetic relationship between the two traits is provided in Supplementary File S1. A total of 200 QTNs were assigned to each trait. To reduce the computational time and memory usage during the simulation, we randomly selected 200 SNPs per chromosome from the 303,002 SNPs, ensuring that all QTNs were included. These 4,000 SNPs (200 SNPs × 20 chromosomes) were used in the subsequent GP model.

**Figure 1.**
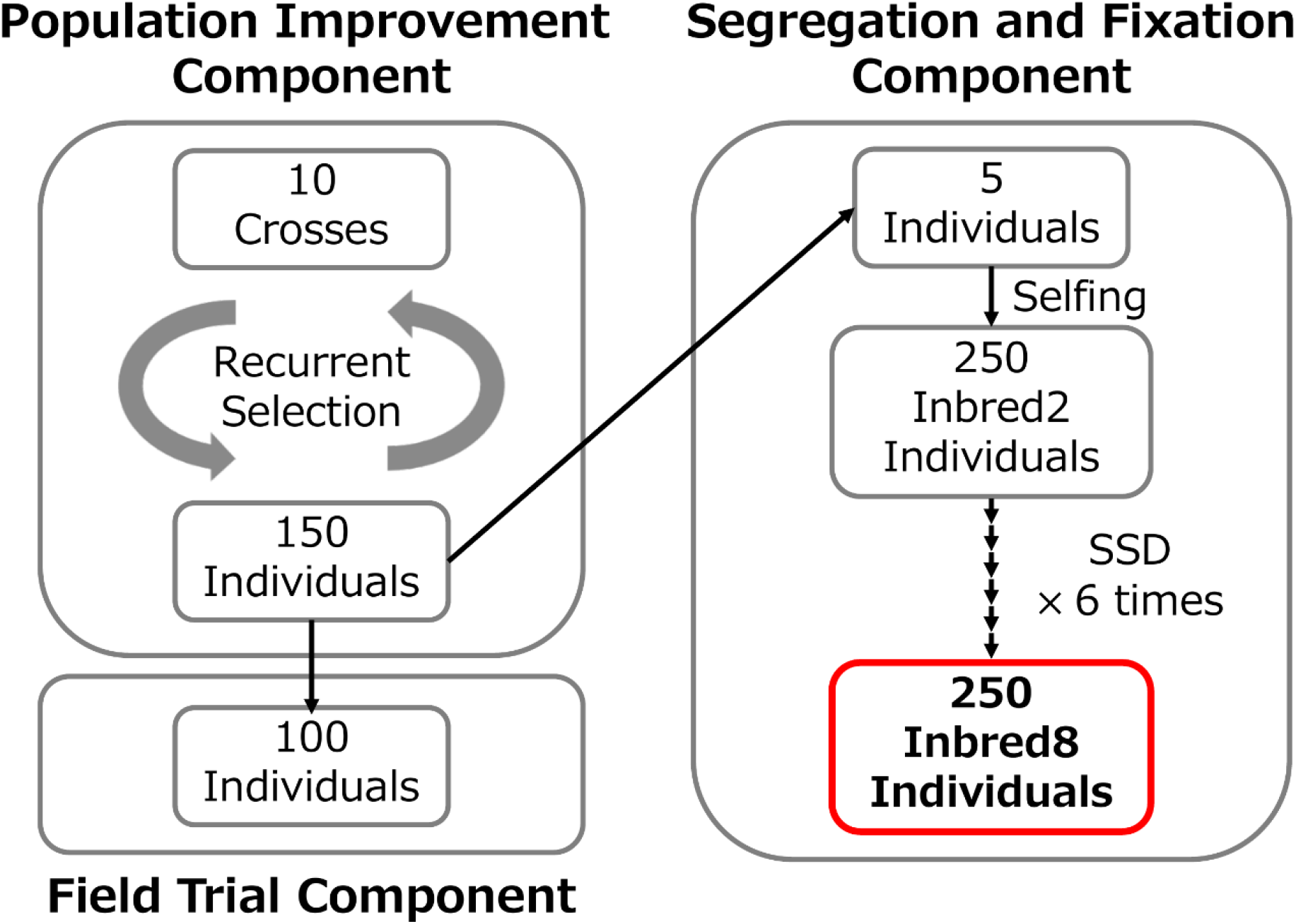
Overview of a breeding program adopted in this breeding simulation. SSD: Single-Seed Descent.

### Phenotypic values

For each simulation, the phenotypic values of the 150 accessions in the initial population were simulated using the following equation:

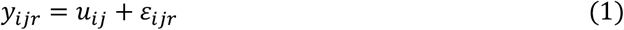

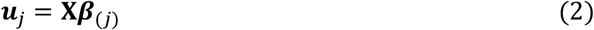

Here, *y*_*ijr*_ represents the phenotypic value of individual *i* (*i* = 1, …, *N*) for trait *j* (*j* = 1, 2) in replicate *r* (*r* = 1, …, *n*_*rep*_, where *n*_*rep*_ = 5), where *N* is the number of individuals in the initial population (*N* = 150), *u*_*ij*_ is the true genotypic value, and *ε*_*ijr*_ is the error value. **X** is the *N* × 4,000 SNP score matrix for the initial population, and ***β***_(*j*)_ is a 4,000 × 1 vector of true SNP effects for trait *j*. The error vector 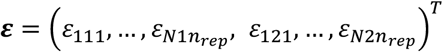 follows a multivariate normal distribution ***ε*** ∼ *MVN*(**0, R**⨂**I**), where **R** is a 2 × 2 residual variance-covariance matrix and **I** is a (*N* × *n*_*rep*_) × (*N* × *n*_*rep*_) identity matrix. The residual variance-covariance matrix **R** is defined as follows:

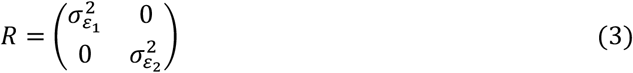

Here, 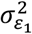 and 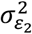 represent the residual variances for Trait1 and Trait2, respectively. For simplicity, we assumed no residual covariance between two traits. The residual variance 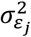 (*j* = 1, 2), was determined using the following equation:

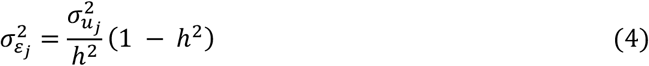

Here, 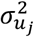 is the genetic variance for trait *j* in the initial population, and *h*^2^ is the individual-based heritability. In this study, we assumed *h*^2^ = 0.6 for both traits. The genetic variance 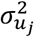 was calculated as follows:

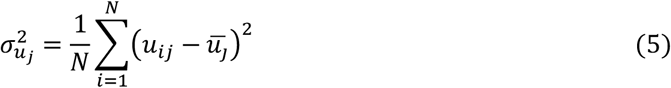

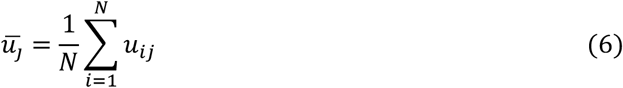

By following the procedure described in Eq. 1–6, the phenotypic values of the initial population can be simulated according to any given heritability.

### Genomic prediction model

To estimate the marker effects of the 4,000 SNPs, a GBLUP (Genomic Best Linear Unbiased Predictor) model was constructed for each trait using the phenotypic values of 150 individuals from the initial population (VanRaden, 2008). The GBLUP model for each trait was expressed as follows:

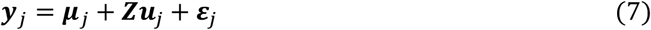

Here, ***y***_*j*_ is a 5*N* × 1 phenotypic value vector for trait *j* (with five replications), ***μ***_*j*_ is the overall mean for trait *j*, ***Z*** is a 5*N* × *N* design matrix, ***u***_*j*_ is an *N* × 1 vector of genotypic values for trait *j*, and ***ε***_*j*_ is a 5*N* × 1 residual vector. The covariance structures are defined as *Var*[***u***_*j*_] ∝ **K** and *Var*[***ε***_*j*_] ∝ **I**, where **K** is an *N* × *N* additive genetic relationship matrix and **I** is an *N* × *N* identity matrix. The matrix **K** was calculated using the *N* × 4,000 SNP score matrix (**X**) of the initial population using the A.mat function in the R package rrBLUP v4.6.3 (Endelman 2011). The GBLUP model was constructed using the EMM.cpp function in the R package RAINBOWR v0.1.38 (Hamazaki and Iwata 2020). Based on the GEBVs 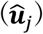 of each trait, the SNP effects for each trait were computed according to the following equation (Strandén and Garrick 2009).

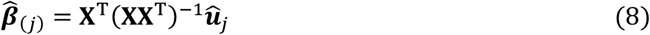

In this equation, 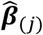 represents the estimated 4,000 × 1 vector of SNP effects for trait *j*, **X** is the *N* × 4,000 SNP score matrix for the initial population, and 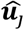 is the *N* × 1 GEBV vector of the initial population. The marker effects in the initial population were estimated using Eq.8 and were used to select individuals and crossing pairs in each breeding strategy. As described later, the GP model was updated throughout the breeding program.

### Breeding program

In this study, we adopted a two-part breeding strategy, originally developed for crops with doubled haploid (DH) protocols (Gaynor et al. 2017), and extended it for use in inbred crops lacking established DH production protocols (Sakurai et al. 2024). We developed and adopted a three-part breeding program by extending the two-part strategy and incorporating an additional Field Trial Component to update the GP model (Figure 2). The Population Improvement Component was aimed at improving the overall genotypic value of the breeding population through rapid recurrent selection. In each cycle, 10 crossing pairs were selected from 150 individuals (with an initial population consisting of 150 accessions). Each pair produced 15 offspring, resulting in 150 progeny individuals (10 pairs × 15 offspring) for the next cycle. In subsequent cycles, 10 new crossing pairs were selected from the 150 newly produced individuals, and 150 offspring were again produced. This process of recurrent selection among highly heterozygous individuals aimed to improve the genotypic value of Trait1 within the population.

**Figure 2.**
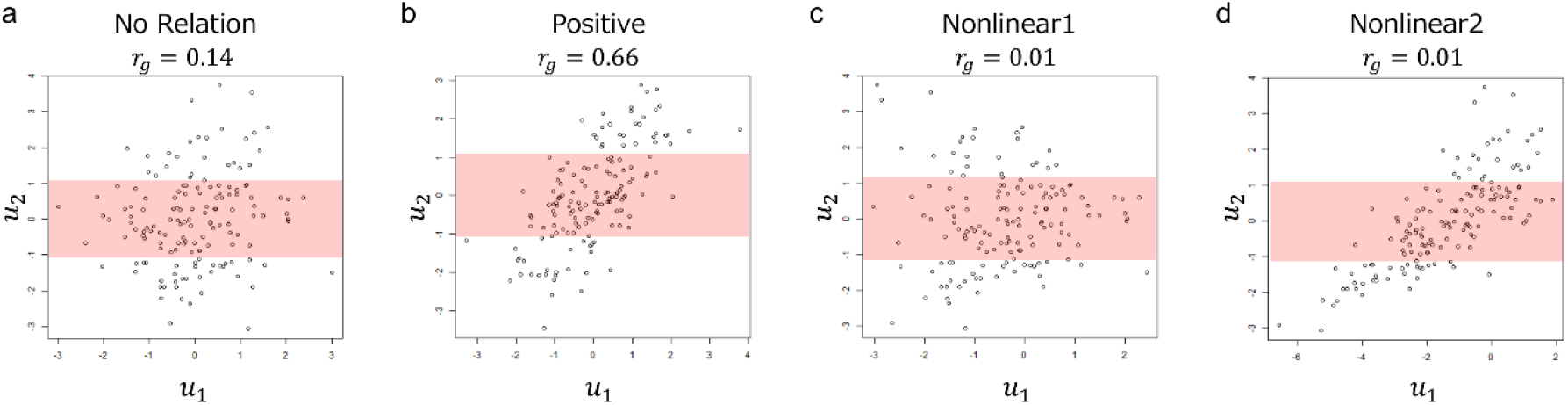
Desirable range of genotypic values for Trait2. Scatterplots of genotypic values produced from the first breeding simulation, in which the relationship between the two traits is attributed to pleiotropy. The red area indicates the desirable range of genotypic values for Trait2. a: No relationship, b: Positive correlation, c: Nonlinear1, d: Nonlinear2. *r*_*g*_: Pearson correlation coefficient between *u*_1_ and *u*_2_.

In soybeans, the number of flowers and the amount of pollen available per plant are limited. Therefore, the maximum number of crossing pairs per individual was set at two. The first generation of 150 individuals produced from 150 accessions consisted of 10 types of *F*_1_ lines. To avoid a rapid decline in genetic diversity within the population, each *F*_1_ line was treated as an individual in the selection of the next 10 crossing pairs, and the maximum number of crossing pairs per *F*_1_ line was limited to two. Additionally, because soybeans produce multiple flowers and can serve as both pollen and seed parents, no restrictions were placed on parental roles with respect to sex. The number of days to seed maturity in ‘Enrei,’ one of Japan’s major soybean cultivars, ranges from 102 to 132 days (Yamada et al. 2012). Based on this and assuming the use of a greenhouse, we considered two cycles per year to be feasible. In this study, we assumed that recurrent selection could be conducted twice a year.

The Segregation and Fixation Component aims to produce new varieties through repeated selfings (Figure 2). Each year (corresponding to even-numbered cycles in the Population Improvement Component), five individuals were selected from the 150 individuals in the Population Improvement Component and sent to the Segregation and Fixation Component for variety development. First, each selected individual underwent selfing to segregate the alleles, producing 50 offspring per individual, resulting in 250 Inbred2 individuals (5 individuals × 50 offspring). These Inbred2 individuals then underwent six rounds of selfings by single seed descent (SSD) method to fix alleles, ultimately yielding 250 Inbred8 individuals per year. In this study, the Inbred2 and Inbred8 generations refer to the individuals derived from one and seven selfings, respectively.

The objective of this breeding program is to produce Inbred8 individuals that exhibit high genotypic values for Trait1 and desirable genotypic values for Trait2 (*l* ≤ *u*_2_ ≤ *h*) for producing a new variety (Figure 2). Because genotypic values for Trait2 in the initial population were generated according to a normal distribution, the desirable range of genotypic values for Trait2 was defined based on the average variance of Trait2 in the initial 150 accessions across all breeding simulations 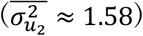. The desirable range was set to include approximately 50% of the initial population, corresponding to the interval −1.08 ≤ *u*_2_ ≤ 1.08 (Figure 1).

In the Field Trial Component, the GP model is updated by conducting field trials to collect phenotypic data. In the conventional two-part strategy, field trials are conducted using doubled haploids (DHs) produced in the Product Development Component (referred to as the Segregation and Fixation Component in this study), and the GP model is updated (Gaynor et al. 2017). However, the use of DHs to update the GP model has two main drawbacks. First, it takes time to produce DHs; however, recurrent selection cycles continue during this period. This time difference resulted in a weaker genetic relationship between the DH population used in the field trials and the breeding population in the Population Improvement Component. This leads to lower prediction accuracy for the GP model. Second, we have to collect genotypic data for all individuals in the DH population used in the field trials. Therefore, additional genotyping costs were incurred. The Field Trial Component in this study was designed to address these issues. Each year, field trials were conducted (in odd-numbered cycles of the Population Improvement Component, excluding the first cycle) using 100 individuals randomly selected from the remaining candidates that were not selected as part of the 10 crossing pairs. Because the genotypic data of these 100 individuals were already collected to select 10 crossing pairs, no additional genotyping costs were incurred. The phenotypic value vector for trait *j* for the newly collected 100 individuals was generated using the following equation:

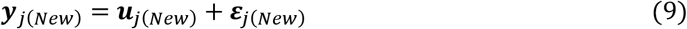

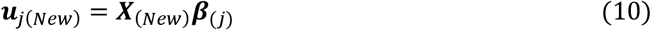

Here, ***y***_*j*(*New*)_ is a 100 × 1 phenotypic value vector for trait *j* of the newly collected 100 individuals, ***u***_*j*(*New*)_ is a 100 × 1 vector of genotypic values, and ***ε***_*j*(*New*)_ is a 100 × 1 error vector. ***X***_(*New*)_ is a 100 × 4,000 SNP score matrix, and ***β***_(*j*)_ is a 4,000 × 1 vector of true SNP effects. In cases where the relationship between the two traits was nonlinear, the values of ***u***_*j*(*New*)_ were updated according to Eq.S5 or S6 in Supplementary File S1. Although genotype-by-year effects are likely to exist in actual field trials, for simplicity, this study did not assume such interactions. The error vector ***ε***_*j*(*New*)_ is generated from a multivariate normal distribution 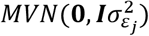, where 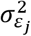 is equivalent to the value defined in Eq.4. In other words, the residual variance is fixed at the value set for the initial population.

When new phenotypic data are collected, 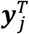 and ***X*** are updated as 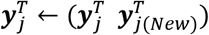 and 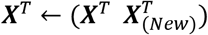 respectively. Using the updated 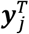 and ***X***, the estimated SNP effect vector 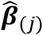 is re-estimated based on Eq.7,8.

The GP models updated through the Field Trial Component were then utilized for selection in subsequent cycles. As the genotypic data used to update the GP models included those of the sib parents of the 150 individuals to be evaluated in the next cycle, an improvement in the prediction accuracy of the GP models was expected. A flowchart of the breeding program is shown in Figure 3.

**Figure 3.**
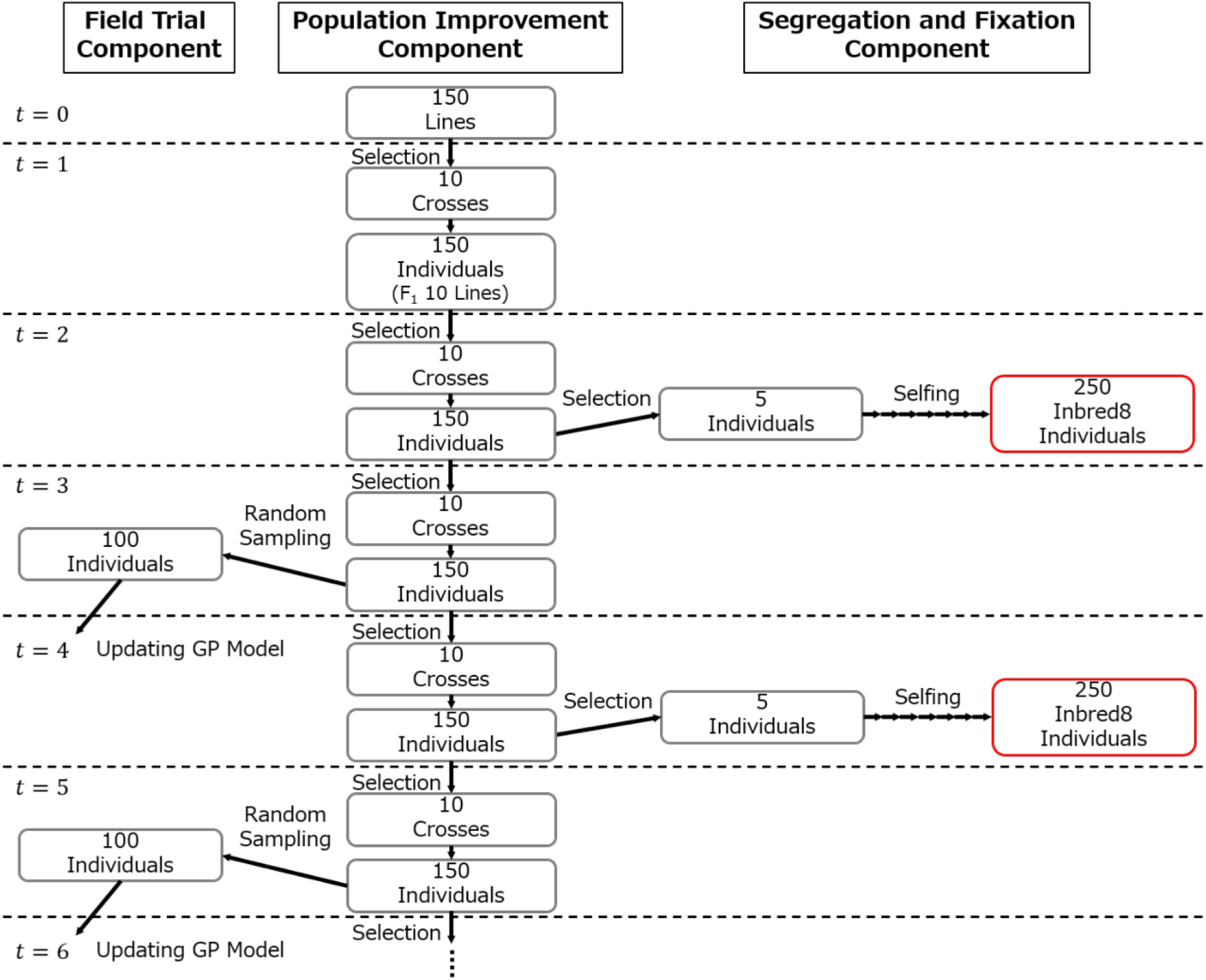
Breeding program flowchart. *t* represents the cycle in the Population Improvement Component. In even-numbered cycles, five individuals were selected for the Segregation and Fixation Component. In odd-numbered cycles, 100 individuals were randomly sampled, and their phenotypic data were collected in the Field Trial Component.

The outcome of a breeding program depends largely on the selection and crossing strategies used in the Population Improvement Component (Figure 2). To evaluate the effectiveness of each breeding strategy, a 10-year breeding program was simulated, and two breeding strategies, Penalized Cross Selection (PCS) and Cross Potential Selection for Multiple Traits (CPS-MT), were compared based on the true genotypic values of the Inbred8 generation produced each year. At each generation, only those individuals among the 250 Inbred8 individuals whose genotypic values for Trait2 were within the desirable range (*l* ≤ *u*_2_ ≤ *h*) were used for evaluation. Over the 10-year breeding program, a total of *T* = 20 cycles of recurrent selection were conducted in the Population Improvement Component. The time required to produce 250 Inbred8 individuals was not considered in the simulation.

### Penalized Cross Selection (PCS)

In PCS, the evaluation value was computed for each crossing pair based on the mean GEBVs for Trait1 and Trait2. Genotypic value for Trait1 should be maximized while that for Trait2 should be in the desirable range (*l* ≤ *u*_2_ ≤ *h*). In PCS, the following value was used as the evaluation value for each crossing pair in the Population Improvement Component:

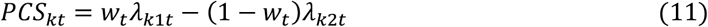

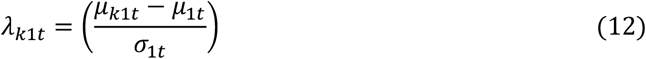

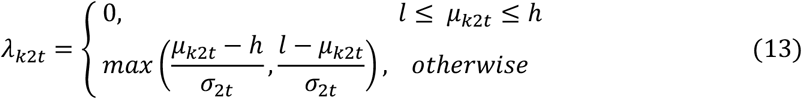

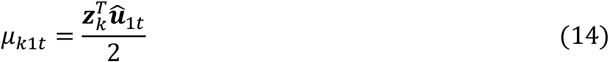

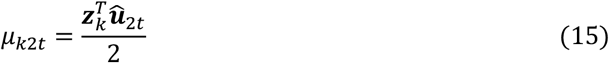

Here, *PCS*_*kt*_ is the evaluation value for crossing pair *k* (*k* = 1, …, *nc*) in cycle *t* (*t* = 1,2, …, *T*, where *T* = 20), and *nc* is the total number of possible crossing pairs 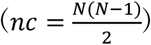, where *N* is the number of individuals in the Population Improvement Component (*N* = 150). *λ*_*k*1*t*_ is the standardized mean GEBV for Trait1 and *λ*_*k*2*t*_ is the penalty value for Trait2 of crossing pair *k. μ*_1*t*_ is the overall mean GEBV 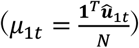 for Trait1 and *σ*_*jt*_ is the standard deviation 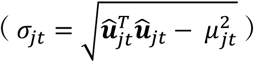 for trait *j* (*j* = 1,2) of the Population Improvement Component, where 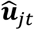 is the *N* × 1 GEBV vector. *μ*_*kjt*_ is the mean GEBV for trait *j* of crossing pair *k*. *w*_*t*_ is a weight parameter. *z*_*k*_ is an *N* × 1 vector, indicating that the two individuals comprise a crossing pair *k*, with ones for those two individuals and zeros elsewhere. The second term in Eq.11 represents a penalty term proportional to the distance from the desirable range for Trait2 (*l* ≤ *u*_2_ ≤ *h*). This encourages selection within the desirable range. PCS is conceptually related to the selection index framework (Hazel and Lush 1942; Hazel 1943), in that it evaluates crossing pairs based on a weighted linear combination of mean GEBVs across traits. Unlike UC, PCS does not incorporate the distribution of progeny GEBVs, and therefore does not evaluate the expected value of the top proportion of progeny. We assumed that *w*_*t*_ should be changed based on the remaining cycles of the breeding program. Nine types of linear trajectories were considered:

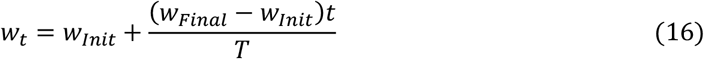

Here, *w*_*t*_ is a weight parameter for cycle *t* (*t* = 1,2, …, *T*) and, *w*_*Init*_ and *w*_*Final*_ are the weight parameters for the first and final cycles. We evaluated all 9 combinations of *w*_*Init*_ ∈ {0.1, 0.5, 0.9} and *w*_*Final*_ ∈ {0.1, 0.5, 0.9}. The process used to determine *w*_*Init*_ and *w*_*Final*_ is described later. PCS selects 10 crossing pairs that maximize the sum of *PCS*_*kt*_ values. The objective is to maximize the following function:

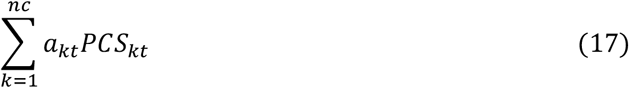

Here, *a*_*kt*_ is a dummy variable (*a*_*kt*_ = 0 if crossing pair *k* is not selected, and *a*_*kt*_ = 1 if it is selected). In this breeding program, the total number of selected crossing pairs was set to 10, and each individual was used in a maximum of two crossing pairs. These constraints are expressed as follows:

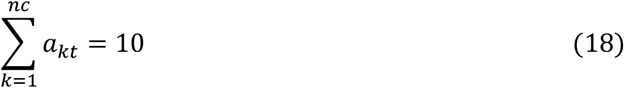

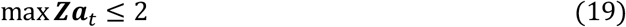

Eq.18 represents the constraint on the total number of selected crossing pairs, while Eq.19 represents the constraint on the maximum number of crossing pairs per individual. Here, ***Z*** is an *N* × *nc* design matrix, defined as ***Z*** = (*z*_1_, …, *z*_*nc*_). Therefore, ***Za***_*t*_ is an *N* × 1 vector that indicates the number of times each individual is used in crossing pairs. In PCS, the optimization problem is to maximize Eq.17 subject to the constraints given in Eq.18,19. Because this optimization problem is an integer programming problem, we can obtain an optimal solution. The “lp” function from the R package “lpSolve” v5.6.23 was used to solve this optimization problem (Berkelaar, 2024). This optimization framework enabled the selection of an optimal set of 10 crossing pairs under PCS in each cycle.

For the values of *w*_*Init*_ and *w*_*Final*_ in Eq.16, a grid search was conducted using all nine combinations across all seven scenarios. For the grid search, 30 independent simulations were performed for each parameter setting. The genetic gain for each generation of Inbred8 individuals was calculated according to Allier et al. (2019a), as follows:

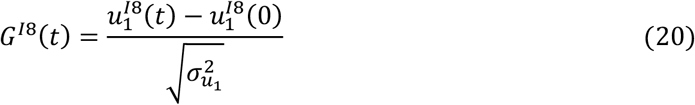

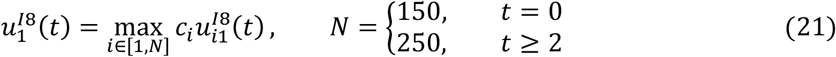

Here 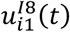 is the true genotypic value for Trait1 of individual *i* in the Inbred8 population produced in cycle (*t* = 2,4, …,20). *c*_*i*_ is a binary variable indicating whether the true genotypic value for Trait2 of individual *i* falls within the desirable range (i.e., *l* ≤ *u*_2_ ≤ *h*); specifically, *c*_*i*_ = 1 if 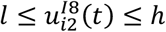, and *c*_*i*_ = 0 otherwise. 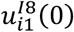 represents the true genotypic value for Trait1 of individual *i* in the initial population of 150 individuals. 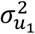 is the genetic variance for Trait1 in the initial population, calculated from the 150 individuals based on Eq.5 in each independent simulation.

For each simulation, the cumulative genetic gain of the Inbred8 population was calculated as follows:

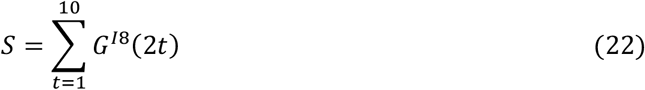

For each of the seven scenarios, the weight parameters (*w*_*Init*_ and *w*_*Final*_) were set to the values that yielded the highest average *S* across the 30 independent simulations conducted for all nine combinations.

### Cross Potential Selection for Multiple Traits (CPS-MT)

CPS-MT is a multi-trait cross selection strategy formulated to handle one target trait and any number of essential traits. The general formulation of CPS-MT for an arbitrary number of essential traits is provided in Supplementary File S1. For ease of presentation, we describe the method for the two-trait case (***J*** = 2), with one target trait (Trait1) and one essential trait (Trait2), which corresponds to the setting used in the present study. CPS-MT selects crossing pairs by considering the progeny segregation in future generations. The genetic covariance between trait *j*_1_ (*j*_1_ = 1,2) and trait *j*_2_ (*j*_2_ = 1,2) in the progeny population after seven rounds of selfings derived from crossing pair *k* in cycle *t*, denoted as 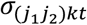, is calculated following (Allier et al. 2019a),

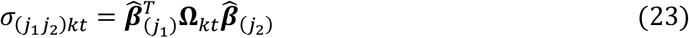

Here, 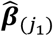 and 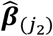 are the 4000 × 1 estimated marker effect vectors for *j*_1_ and *j*_2_, and **Ω**_*kt*_ represents the 4,000 × 4,000 marker variance-covariance matrix of the progeny population after seven rounds of selfings. The detailed calculation procedure of **Ω**_*kt*_ is provided in Supplementary File S1.

In CPS-MT, the GEBV distribution for the Inbred8 population was utilized, and each crossing pair was evaluated based on the genetic potential of the superior genotypes that could be produced in the Inbred8 generation. We assume that the GEBV distribution of the Inbred8 population derived from crossing pair *k* follows the following multivariate normal distribution, calculated using 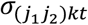 obtained from Eq.23, and *μ*_*k*1*t*_ and *μ*_*k*2*t*_ obtained from Eq.14,15.

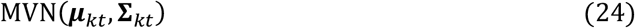

Here, ***μ***_*kt*_ = (*μ*_*k*1*t*_ *μ*_*k*2*t*_)^*T*^ is the 2 × 1 vector of mean GEBVs, and **∑**_*kt*_ is the 2 × 2 genetic variance-covariance matrix between traits, where the (*j*_1_, *j*_2_) element corresponds to 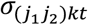. In PCS, only ***μ***_*kt*_ was used to evaluate each crossing pair, whereas CPS-MT evaluates each crossing pair by utilizing both ***μ***_*kt*_ and **∑**_*kt*_. In the breeding program, 15 progeny individuals were produced from each selected crossing pair for the Population Improvement Component. The individuals selected for the Segregation and Fixation Component underwent repeated selfings to produce 50 Inbred8 individuals. Therefore, when evaluating each cross pair, the expected evaluation value assumed that 750 Inbred8 individuals (15 progeny × 50 Inbred8 individuals) were produced. We simulated 750 samples from the predicted progeny distribution (Eq.24) and computed the same evaluation value in Eq.11 for each sample as follows:

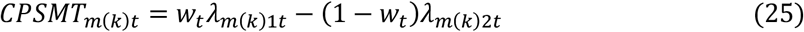

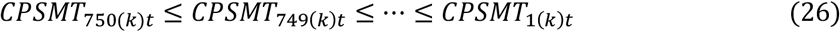

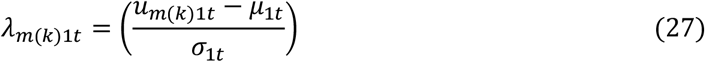

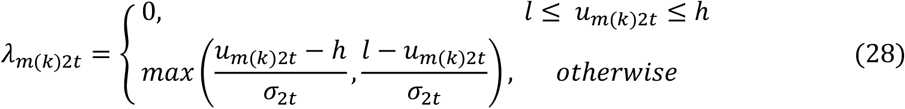

Here, *CPSMT*_*m*(*k*)*t*_ is the ordered evaluation value for sample *m* (*m* = 1, …, 750), *u*_*m*(*k*)*jt*_ is the simulated value from Eq.24 (i.e. ***u***_*m*_(_*k*_)_*t*_ ∼ MVN(***μ***_*kt*_, **∑**_*kt*_)). *w*_*t*_, *μ*_1*t*_ and *σ*_*jt*_ are the same as in Eq.11. Each crossing pair was evaluated based on a UC-like value as follows:

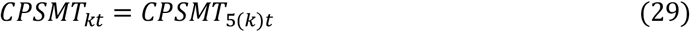

Here *CPSMT*_*kt*_ is the evaluation value for cross *k*. UC quantifies the expected value of superior individuals derived from a given cross, whereas CPS-MT directly simulates progenies and evaluates a given cross based on the simulated superior individuals. In this breeding program, the top five individuals were treated as superior individuals because top five individuals will be selected for the Segregation and Fixation Component (Figure 1). Unlike PCS, which evaluates crossing pairs based solely on the mean GEBVs, CPS-MT explicitly accounts for the GEBV distribution of the Inbred8 progeny derived from each crossing pair, thereby incorporating the genetic potential of parental combinations beyond their mean performance (Figure 4). This selection algorithm can also be extended to multi-trait constraints. The details are provided in Supplementary File S1. We repeated this sampling simulation 20 times and used the mean value of *CPSMT*_*kt*_ to obtain a stable estimate of *CPSMT*_*kt*_.

**Figure 4.**
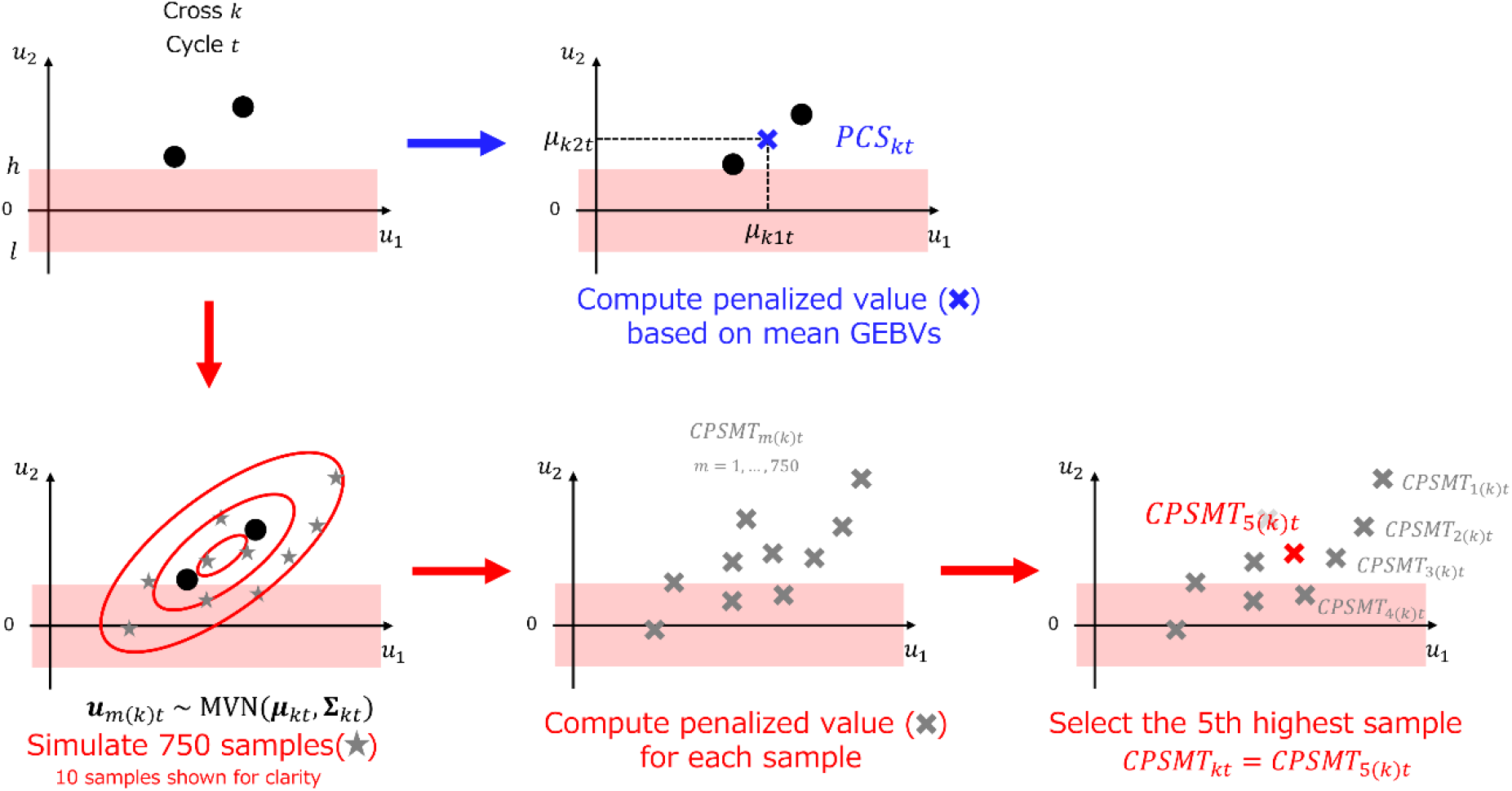
Schematic overview of the evaluation procedures for PCS and CPS-MT. Top row (blue): In Penalized cross selection (PCS), the evaluation value *PCS*_*kt*_ for crossing pair *k* in cycle *t* is computed as a penalized value based on the mean GEBVs of the crossing pair. Bottom row (red): In Cross-Potential Selection for Multiple Traits (CPS-MT), 750 samples were simulated from the predicted progeny distribution (Eq.24; 10 samples are shown for clarity). The penalized value *CPSMT*_*m*_(_*k*_)_*t*_ is computed for each sample *m*, and the samples were ranked in descending order. The evaluation value for crossing pair *k* is defined as the 5^th^ highest value, *CPSMT*_*kt*_ = *CPSMT*_5_(_*k*_)_*t*_. ***μ***_*kt*_: Mean GEBV vector, **∑**_*kt*_: Genetic variance-covariance matrix between traits.

In CPS-MT, 10 crossing pairs were selected to maximize the sum of the *CPSMT*_*kt*_ values. The objective function is defined as

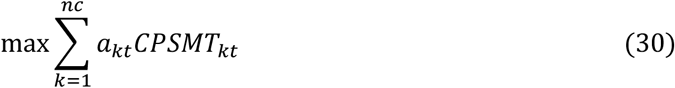

Here, *a*_*kt*_ is a dummy variable for each crossing pair *k* (*a*_*kt*_ = 0 indicates that the crossing pair *k* is not selected, whereas *a*_*kt*_ = 1 indicates that it is selected). Because each individual can be used for at most two crossing pairs and the number of crossing pairs to be selected is 10, CPS-MT maximizes Eq.30 under the same constraints as in PCS, namely Eq.18,19. This optimization problem is also a type of integer programming problem and was solved using the “lp” function from the R package “lpSolve” v5.6.23 (Berkelaar, 2024). Thus, an optimal set of 10 crossing pairs was selected for each cycle using CPS-MT.

As in PCS, the weight parameter *w*_*t*_ in CPS-MT was determined by grid search over all 9 combinations of *w*_*Init*_ ∈ {0.1, 0.5, 0.9} and *w*_*Final*_ ∈ {0.1, 0.5, 0.9}.

### Selection of individuals for the Segregation and Fixation Component

In the Segregation and Fixation Component, five individuals were selected each year from the 150 individuals in the Population Improvement Component. These five individuals were selected using a procedure based on *CPSMT*_*kt*_, which was applied consistently across the two strategies. Specifically, for each individual *i*, ***μ***_*it*_ and ***∑***_*it*_ in Eq.24 were calculated to estimate the GEBV distribution of the Inbred8 population that would be produced from individual *i* through seven rounds of selfings. At this point, by regarding crossing pair *k* as consisting of individual *i* with itself and setting the number of selfing rounds to six, the procedure can be naturally extended by treating selfing as a special case of crossing. Because 50 Inbred8 individuals will be produced from a selected individual, we simulated 50 samples following MVN(***μ***_*it*_, **∑**_*it*_) and *CPSMT*_*m*_(_*i*_)_*t*_ (*m* = 1, …, 50) was computed based on the Eq.25 – 28. We evaluated individuals as follows:

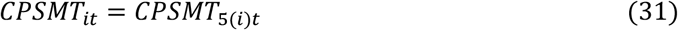

Here, *CPSMT*_*it*_ is the evaluation value for individual *i*. The top five individuals (top 10%) were treated as superior individuals. We repeated this sampling simulation 200 times and used the mean value of *CPSMT*_*it*_ to obtain a stable estimate of *CPSMT*_*it*_.

The five individuals with the highest *CPSMT*_*it*_ values were selected for the Segregation and Fixation Component. The selected individuals were also allowed to be used as crossing pairs in the Population Improvement Component.

### Comparison

After completing the grid search for the parameter *w*_*t*_ across all seven scenarios for PCS and CPS-MT, the results of the breeding simulations conducted using the optimized *w*_*t*_ were compared between the two strategies. The genetic gain in the Inbred8 generation was evaluated for each strategy at cycle *t* (*t* = 2, 4, …, 20) using *G*_*I*8_(*t*), which was calculated using Eq.20. In addition, the genetic variance in the Population Improvement Component serves as a source of genetic improvement. Genetic variance in the Population Improvement Component was calculated as follows:

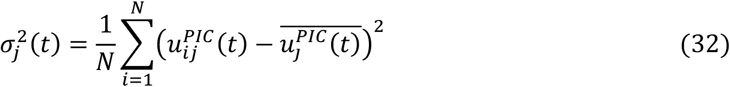

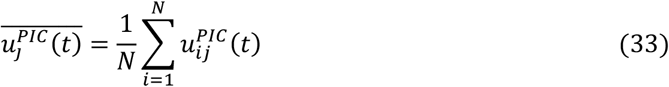

Here 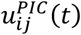 is the true genotypic value for trait *j* (*j* = 1,2) of individual *i* in cycle *t* (*t* = 1,2, …,20) of the Population Improvement Component. The prediction accuracy for trait *j* in each cycle of the Population Improvement Component was calculated as the Pearson correlation coefficient between the true genotypic values and GEBVs of the 150 individuals. Finally, in each cycle of the Population Improvement Component, the prediction accuracies of the genetic (co)variances between trait *j*_1_and trait *j*_2_ (*j*_1_, *j*_2_ = 1,2) in the Inbred8 generation were calculated as the Pearson correlation coefficient between 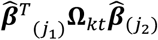 and 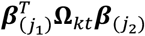. In the cases of “Nonlinear1” and “Nonlinear2,” the genotypic value of Trait1 changes as a function of the genotypic value of Trait2, according to Eq.S5 and S6, respectively. Consequently, the genotypic values in the Inbred8 population no longer followed a multivariate normal distribution. Therefore, the prediction accuracy of the genetic (co)variance between traits in the Inbred8 generation was calculated only for the cases in which the relationship between traits was “No Relation” or “Positive.” The prediction accuracy of genetic (co)variances in the progeny depends on the estimated accuracy of marker effects (Eq.23). In each cycle, the prediction accuracy was evaluated as the Pearson correlation coefficient between the true marker effects (***β***_(*j*)_) and the estimated marker effects 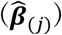, using only the unfixed markers in the population.

## Results

### Determination of the weight parameter and cumulative genetic gain

Tables 1 and 2 present the results of the grid search for the weight parameters *w*_*Init*_ and *w*_*Final*_ in Eq.16 for PCS and CPS-MT, respectively. The value in each cell represents the average cumulative genetic gain calculated using Eq.22, while the numbers in parentheses indicate the number of failed simulations. A failed simulation was defined as a simulation run in which no genotype in the Inbred8 population satisfied the predefined desirable range of genotypic values for Trait2 in at least one generation.

**Table1.**
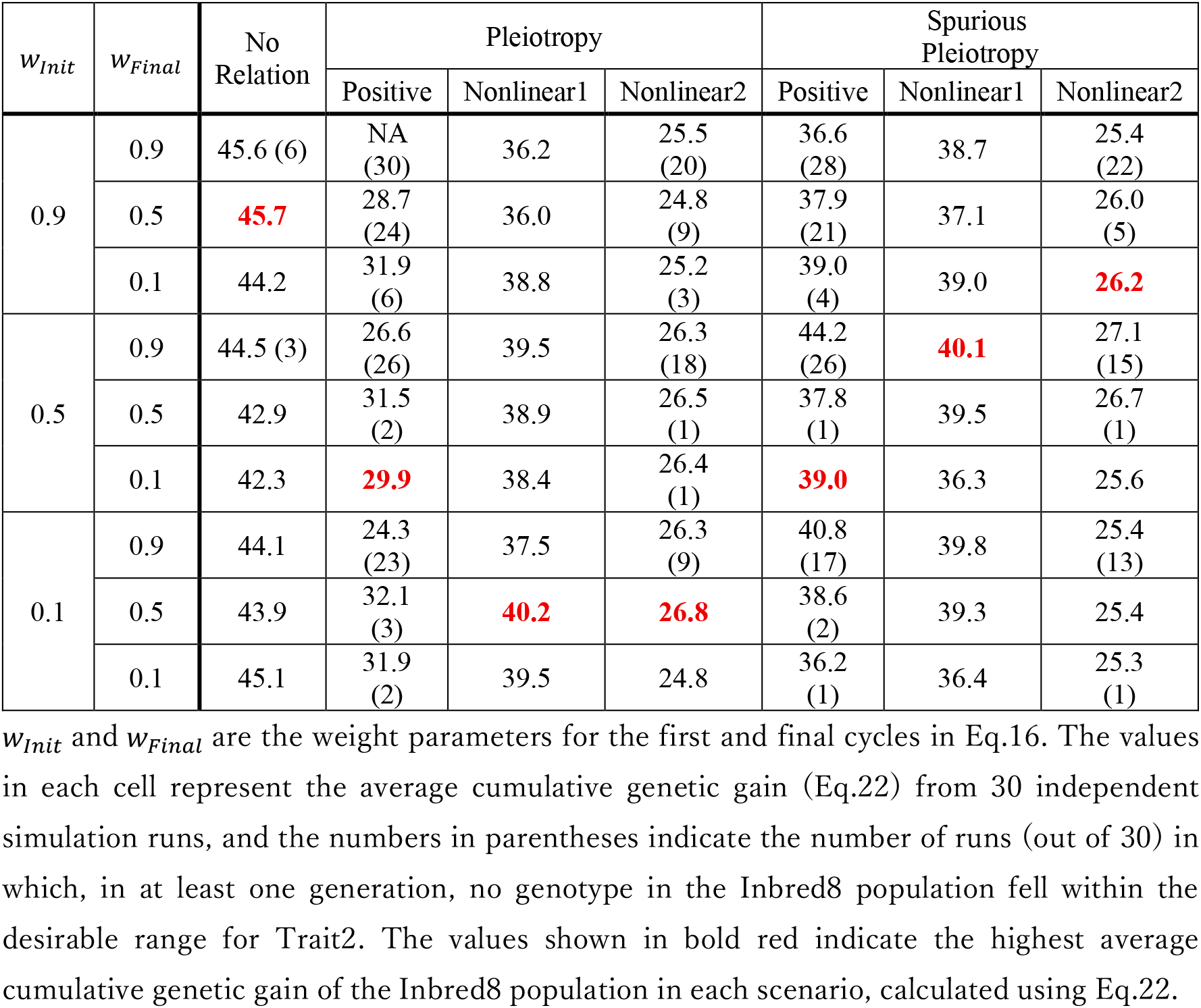
Grid search for the combination of *w*_*Init*_ and *w*_*Final*_ in PCS.

| $w_{Init}$ | $w_{Final}$ | No<br>Relation | Pleiotropy | | | Spurious<br>Pleiotropy | | |
| --- | --- | --- | --- | --- | --- | --- | --- | --- |
|  |  |  | Positive | Nonlinear1 | Nonlinear2 | Positive | Nonlinear1 | Nonlinear2 |
| 0.9 | 0.9 | 45.6 (6) | NA<br>(30) | 36.2 | 25.5<br>(20) | 36.6<br>(28) | 38.7 | 25.4<br>(22) |
|  | 0.5 | **45.7** | 28.7<br>(24) | 36.0 | 24.8<br>(9) | 37.9<br>(21) | 37.1 | 26.0<br>(5) |
|  | 0.1 | 44.2 | 31.9<br>(6) | 38.8 | 25.2<br>(3) | 39.0<br>(4) | 39.0 | **26.2** |
| 0.5 | 0.9 | 44.5 (3) | 26.6<br>(26) | 39.5 | 26.3<br>(18) | 44.2<br>(26) | **40.1** | 27.1<br>(15) |
|  | 0.5 | 42.9 | 31.5<br>(2) | 38.9 | 26.5<br>(1) | 37.8<br>(1) | 39.5 | 26.7<br>(1) |
|  | 0.1 | 42.3 | **29.9** | 38.4 | 26.4<br>(1) | **39.0** | 36.3 | 25.6 |
| 0.1 | 0.9 | 44.1 | 24.3<br>(23) | 37.5 | 26.3<br>(9) | 40.8<br>(17) | 39.8 | 25.4<br>(13) |
|  | 0.5 | 43.9 | 32.1<br>(3) | **40.2** | **26.8** | 38.6<br>(2) | 39.3 | 25.4 |
|  | 0.1 | 45.1 | 31.9<br>(2) | 39.5 | 24.8 | 36.2<br>(1) | 36.4 | 25.3<br>(1) |

$w_{Init}$ and $w_{Final}$ are the weight parameters for the first and final cycles in Eq.16. The values in each cell represent the average cumulative genetic gain (Eq.22) from 30 independent simulation runs, and the numbers in parentheses indicate the number of runs (out of 30) in which, in at least one generation, no genotype in the Inbred8 population fell within the desirable range for Trait2. The values shown in bold red indicate the highest average cumulative genetic gain of the Inbred8 population in each scenario, calculated using Eq.22.

**Table2.** Grid search for the combination of *w*_*Init*_ and *w*_*Final*_ in CPS-MT.

| $w_{Init}$ | $w_{Final}$ | No Relation | Pleiotropy | | | Spurious Pleiotropy | | |
| --- | --- | --- | --- | --- | --- | --- | --- | --- |
|  |  |  | Positive | Nonlinear1 | Nonlinear2 | Positive | Nonlinear1 | Nonlinear2 |
| 0.9 | 0.9 | 48.5 | NA (30) | 42.6 | 27.7 (10) | 46.1 (28) | 43.5 | 27.8 (13) |
|  | 0.5 | 51.1 | 34.2 (22) | 41.2 | 28.4 (3) | 39.1 (15) | 43.8 | 29.1 |
|  | 0.1 | 50.3 | 35.8 (6) | 43.5 | 28.7 | 43.5 (4) | 42.6 | **30.0** |
| 0.5 | 0.9 | 50.9 | 33.9 (23) | 45.1 | 29.4 (7) | 45.3 (16) | **46.5** | 27.4 (11) |
|  | 0.5 | 50.5 | 37.2 (4) | 43.6 | 30.4 (1) | 42.9 (2) | 43.4 | 29.2 |
|  | 0.1 | 50.8 | 33.8 (1) | 43.7 | **29.6** | **43.3** | 43.9 | 28.7 |
| 0.1 | 0.9 | 48.9 | 34.8 (18) | 42.7 | 28.8 | 45.8 (9) | 43.6 | 29.3 (7) |
|  | 0.5 | **51.6** | 35.5 (1) | 43.0 | 28.9 | 44.4 (2) | 42.6 | 29.3 |
|  | 0.1 | 48.2 | **35.5** | **45.4** | 28.6 | 44.1 (1) | 45.1 | 29.4 |

$w_{Init}$ and $w_{Final}$ are the weight parameters for the first and final cycles in Eq.16. The values in each cell represent the average cumulative genetic gain (Eq.22) from 30 independent simulation runs, and the numbers in parentheses indicate the number of runs (out of 30) in which, in at least one generation, no genotype in the Inbred8 population fell within the desirable range for Trait2. The values shown in bold red indicate the highest average cumulative genetic gain of the Inbred8 population in each scenario, calculated using Eq.22.

In both PCS and CPS-MT strategies, when a positive correlation existed between traits, a greater number of simulation failures were observed under large weight parameters (Tables 1 and 2). With a large weight parameter in the final cycle (*w*_*Final*_ = 0.9), the proportion of Inbred8 individuals satisfying the constraint progressively declined across generations in both PCS and CPS-MT (Figures S2 and S3). A similar trend was also observed in the “Nonlinear2” scenario. In the case of “Nonlinear2,” a positive correlation arises between the two traits within the desirable genotypic range for Trait2 (−1.08 ≤ *u*_2_ ≤ 1.08) (Figure 2D). Therefore, this scenario exhibits a tendency similar to that of the positive correlation case. In contrast, under the “Nonlinear1” scenario, desirable Inbred8 individuals could be generated even with relatively large values of *w*_*Init*_ and *w*_*Final*_. This is because, in “Nonlinear1”, genotypes whose genotypic values for Trait2 fall within the desirable range tend to have higher genotypic values for Trait1 (as defined in Eq.S5). Hence, desirable Inbred8 individuals could be produced even when the value of *w*_*t*_ was relatively large. Regardless of the combinations of *w*_*Init*_ and *w*_*Final*_ in Eq.16, CPS-MT consistently achieved a higher cumulative genetic gain than that of PCS across all scenarios, even without accounting for the number of failed simulations (Tables 1 and 2).

In the following analysis, the simulation results were based on the best combination of *w*_*Init*_ and *w*_*Final*_ that yielded the highest average cumulative genetic gain in each scenario. In PCS, the combination of “Spurious Pleiotropy” and “Positive” resulted in a 30% higher cumulative genetic gain compared to the combination of “Pleiotropy” and “Positive”; in CPS-MT, the increase was 22%. These results suggest that when the cause of trait correlation is pleiotropy, it is difficult to improve only one of the traits, regardless of the crossing strategy. On the other hand, in the combination of “Spurious Pleiotropy” and “Nonlinear1,” the increase in cumulative genetic gain compared to “Pleiotropy” and “Nonlinear1” was limited to 2% under CPS-MT. Similarly, the combination of “Spurious Pleiotropy” and “Nonlinear2” showed only a 2% increase compared to the “Pleiotropy” and “Nonlinear2” under CPS-MT. Furthermore, in the nonlinear scenarios, no substantial difference was observed between “Spurious Pleiotropy” and “Pleiotropy” when using PCS.

When comparing PCS and CPS-MT, the cumulative genetic gain achieved using CPS-MT was, on average, 14% higher than that of PCS across all seven scenarios (Tables 1 and 2). Furthermore, in the combination of “Pleiotropy” and “Positive,” CPS-MT showed 19% higher cumulative genetic gain compared to PCS. These results indicate that CPS-MT, which utilizes the distribution of GEBVs in the Inbred8 population, is more effective than PCS in various situations.

### Genetic gain in Inbred8 population

Figure 5 shows the time-series changes in genetic gain of the Inbred8 population. In every scenario, the difference between PCS and CPS-MT increased from the early to later generations, and in the final generation (*t* = 20), the genetic gain of CPS-MT was on average 17% higher than that of PCS. In the combination of “Pleiotropy” and “Positive,” the genetic gain achieved by CPS-MT exceeded that of PCS by an average 17% across all generations. The maximum advantage of PCS over CPS-MT was observed under the combination of “Pleiotropy” and “Nonlinear1” in the 2^nd^ generation (*t* = 2), where PCS exceeded CPS-MT by 2%. However, in the 4^th^ generation (*t* = 4), CPS-MT outperformed PCS by 7%.

**Figure 5.**
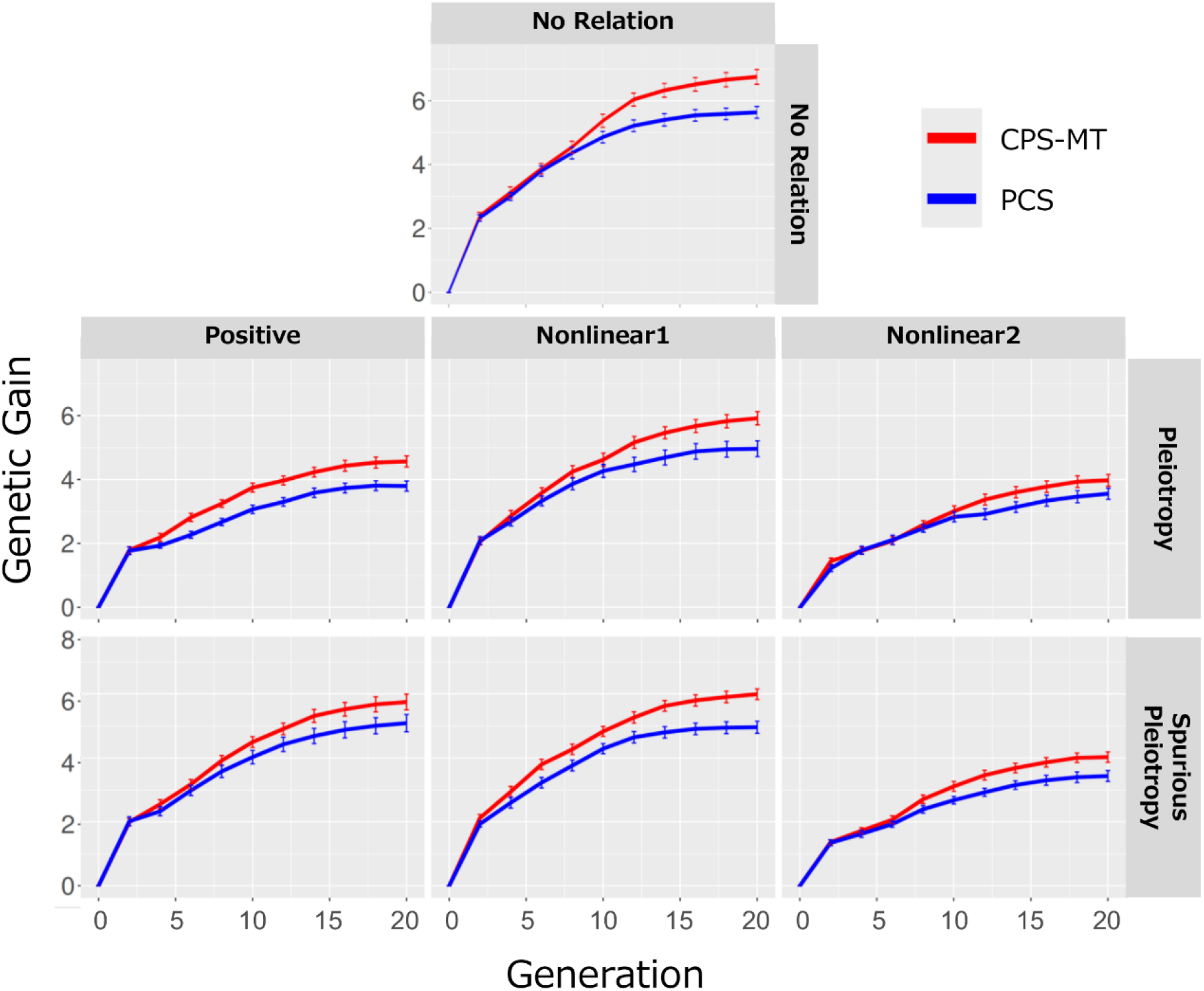
Genetic gain in the Inbred8 population. Time-series changes over 30 independent breeding simulations for each of the seven scenarios. Error bars represent standard error. CPS-MT: Cross Potential Selection for Multiple Traits, PCS: Penalized Cross Selection.

### Genetic variance in population improvement component

Figure 6 shows the time-series changes in the genetic variance of Trait1 in the Population Improvement Component. In every scenario, the genetic variance largely decreases in the 1^st^ generation (*t* = 1) compared to the initial generation (*t* = 0), because 150 individuals are derived from only 10 types of F_1_ lines in the 1^st^ generation. In the 2^nd^ generation, 150 genetically different individuals were produced, leading to the recovery of the genetic variance of Trait1. In all scenarios, the genetic variance for Trait1 tends to decrease as the generations progress, indicating that genetic gain was achieved at the cost of genetic variance. CPS-MT tended to maintain higher genetic variance for Trait1 compared to PCS in “No Relation” and “Nonlinear1” scenarios (Figure 6). From the 7^th^ generation (*t* = 7), CPS-MT consistently retained higher genetic variance than PCS across all scenarios. A similar trend was observed for the genetic variance of Trait2 (Figure S4). In the final generation (*t* = 20), the genetic variance of Trait1 under PCS retained, on average, only 2.6% of the initial population’s genetic variance across all seven scenarios, whereas CPS-MT retained 6.7% (Figure 6). For Trait2, PCS maintained 5.5% and CPS-MT 8.6% of the initial genetic variance, respectively (Figure S4).

**Figure 6.**
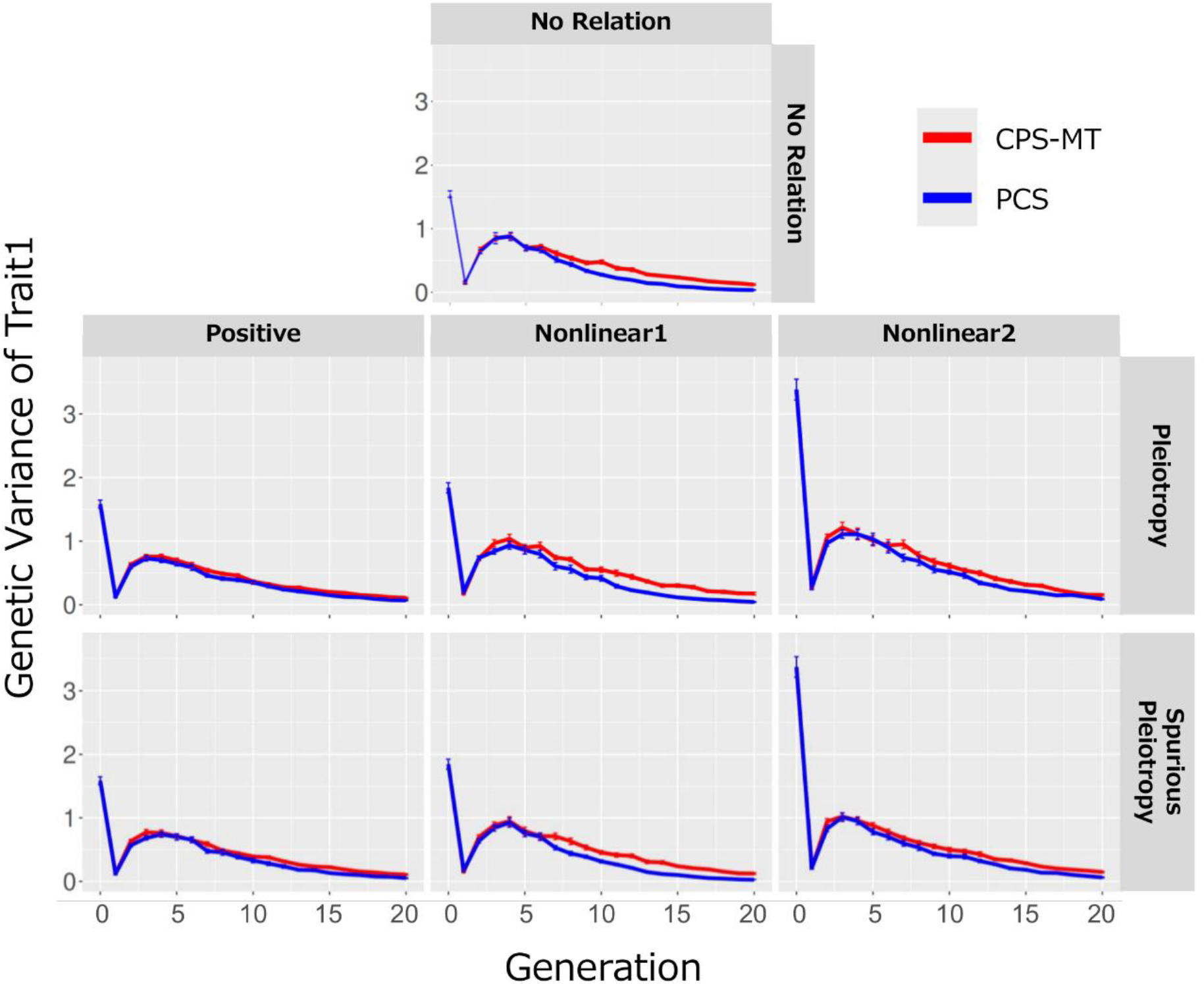
Genetic variance of Trait1 in Population Improvement Component. Time-series changes over 30 independent breeding simulations for each of the seven scenarios. Error bars represent standard error. CPS-MT: Cross Potential Selection for Multiple Traits, PCS: Penalized Cross Selection.

### Prediction accuracy of genotypic value in Population Improvement Component

Figure 7 shows the prediction accuracy of the genotypic value of Trait1 in the Population Improvement Component. Because the GP model was updated in each even-numbered generation from the 4^th^ generation (*t* = 4) using the phenotypic data obtained from the Field Trial Component, the prediction accuracy changed across generations. For Trait1, CPS-MT showed an increasing trend in prediction accuracy after the 10^th^ generation (*t* = 10), whereas PCS exhibited a plateau or even a declining trend (Figure 7). In the final generation (*t* = 20), the prediction accuracy of CPS-MT was, on average, 68% higher than that of PCS across all seven scenarios. In particular, for the scenarios “No Relation” and “Nonlinear1,” the average prediction accuracy in the final generation (*t* = 20) was 0.70 for CPS-MT, compared to 0.31 for PCS across the three relevant scenarios. For Trait2, no substantial difference in prediction accuracy was observed between CPS-MT and PCS in any scenario (Figure S5).

**Figure 7.**
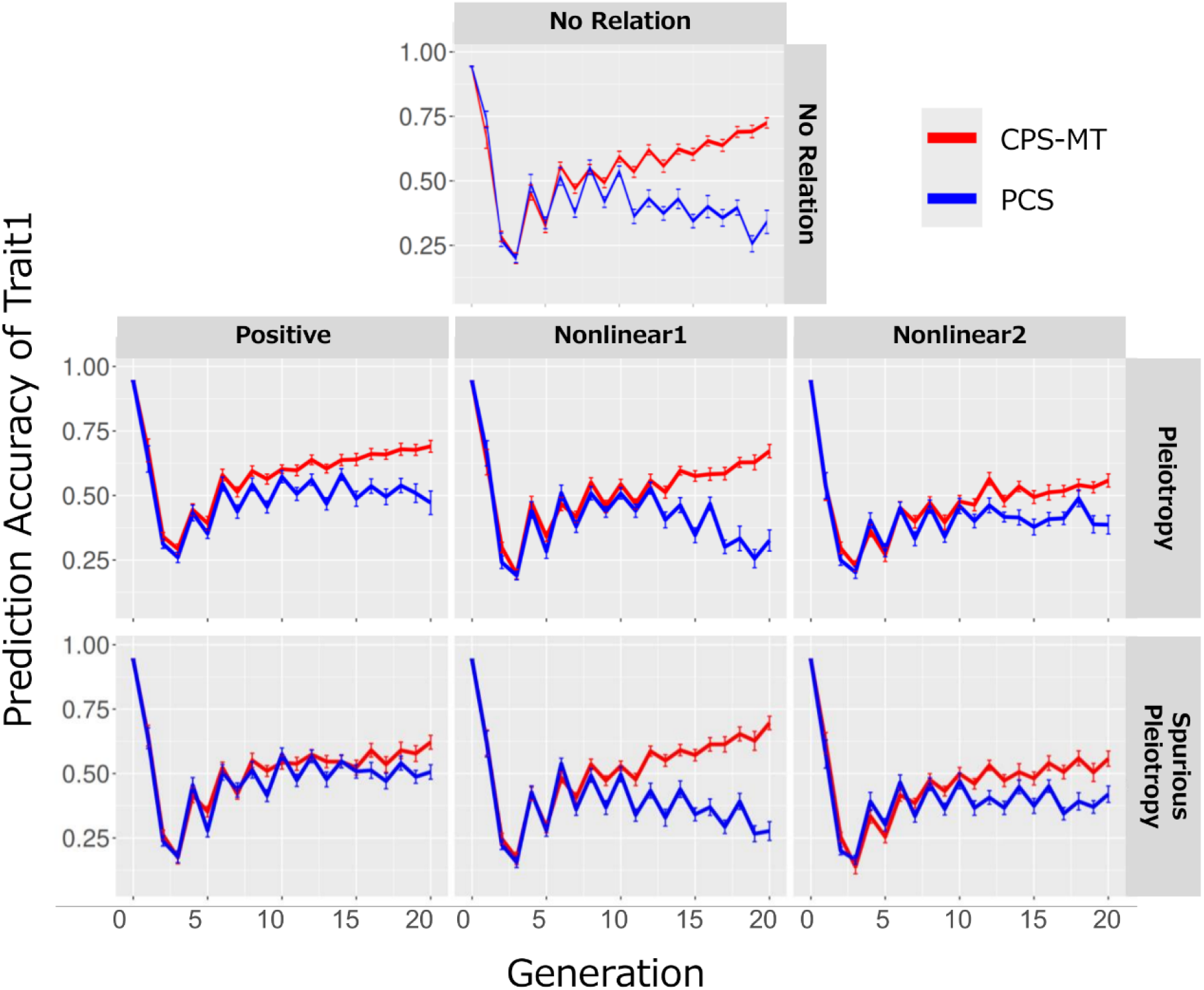
Prediction accuracy of Trait1 in Population Improvement Component. Time-series changes over 30 independent breeding simulations for each of the seven scenarios. Error bars represent standard error. CPS-MT: Cross Potential Selection for Multiple Traits, PCS: Penalized Cross Selection.

### Prediction accuracy of genetic (co)variances in the Inbred8 population

Figure 8 shows the prediction accuracy of the genetic (co)variance in the Inbred8 population. The prediction accuracy of *σ*_(11)_, *σ*_(22)_, and *σ*_(12)_ improved as generations advanced, exhibiting a trend similar to the changes in the prediction accuracy of genotypic values for Trait1 and Trait2 (Figures 7 and S5). The average prediction accuracies of *σ*_(11)_, *σ*_(22)_, and *σ*_(12)_ across the three scenarios in the final generation (*t* = 20) were higher by 149%, 120%, 243%, respectively, than those in the 1^st^ generation (*t* = 1). In all scenarios, the prediction accuracy of the marker effects improved with advancing generations (Figure S6). The mean estimated accuracies of the marker effects for Trait1 and Trait2 increased by 98% and 116%, respectively, from the 1^st^ generation (*t* = 1) to the final generation (*t* = 20) across the three scenarios.

**Figure 8.**
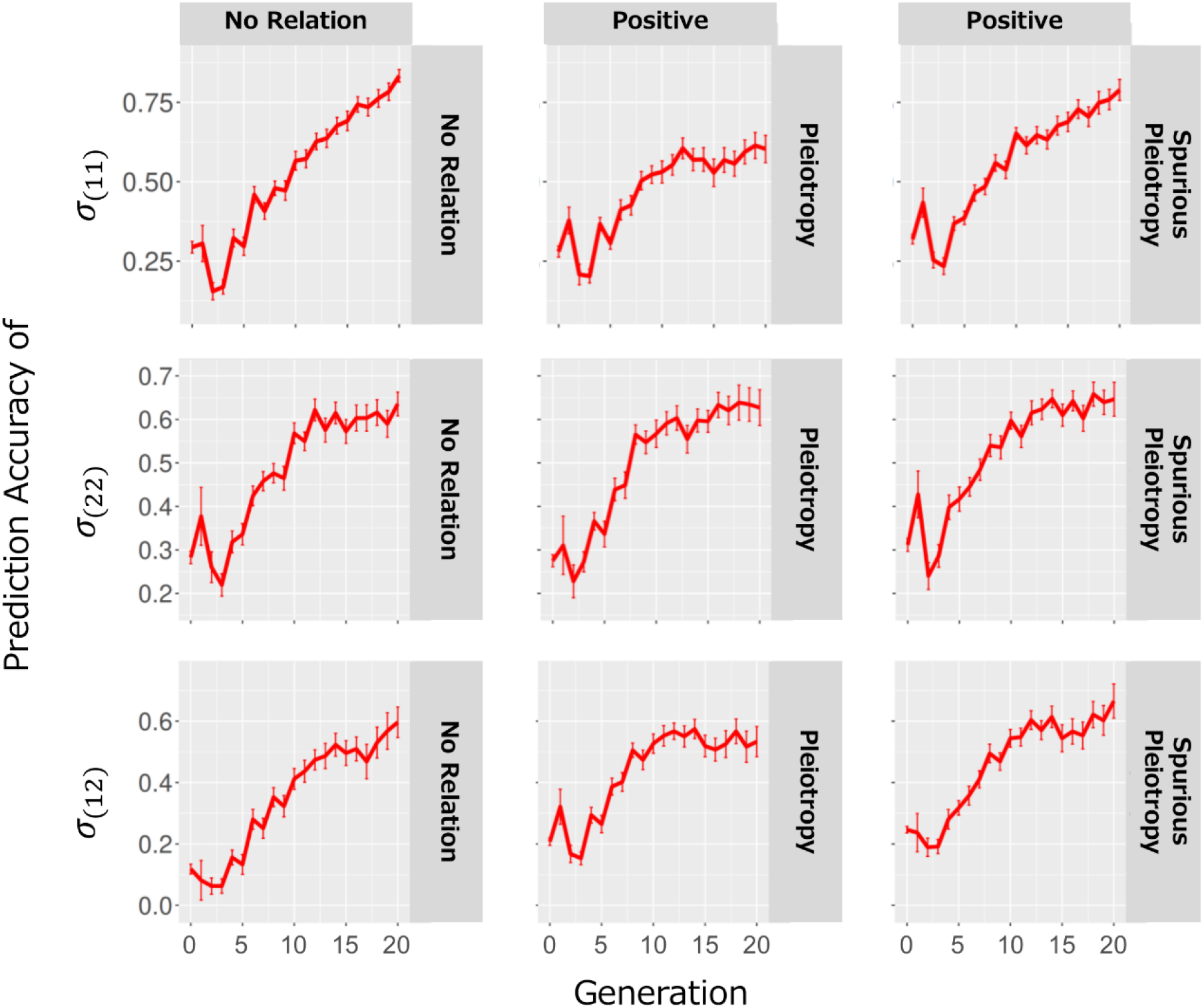
Prediction accuracies of genetic (co)variances for Inbred8 population in CPS-MT. Time-series changes over 30 independent breeding simulations for each of the three scenarios. Error bars represent standard error. CPS-MT: Cross Potential Selection for Multiple Traits, *σ*_(11)_: genetic variance of Trait1, *σ*_(22)_: genetic variance of Trait2, *σ*_(12)_: genetic covariance between Trait1 and Trait2.

## Discussion

### Effectiveness of CPS-MT

In this study, the usefulness of CPS-MT over PCS was demonstrated across all combinations of trait relationships and their underlying causes, particularly in achieving sustained genetic gain while constraining an essential trait within a desirable range (Tables 1, 2, and Figure 5). While PCS selected crossing pairs based solely on their GEBVs, CPS-MT selected crossing pairs considering the GEBV distribution of the progeny (Inbred8 individuals) produced from each crossing pair (Figure 4). These results indicate the importance of considering GEBV distribution in the progeny population for target trait improvement under other trait constraints. Previous studies have also indicated the effectiveness of breeding strategies that utilize UC (Allier et al. 2019b; Allier et al. 2020; Sanchez et al. 2023; Sakurai et al. 2024), the expected value of superior individuals derived from each crossing pair (Schnell and Utz 1975). However, these strategies only considered UC for the target trait and have not accounted for other traits. CPS-MT differs from these approaches in that it explicitly incorporates progeny GEBV distributions under multi-trait constraints. Although breeding strategies utilizing UC to simultaneously improve the genotypic values of the two target traits have been proposed (Atanda and Bandillo 2024; Kinoshita et al. 2025), no previous studies have evaluated the effectiveness of incorporating the distribution of GEBVs into the progeny population. In particular, such an evaluation has not been conducted under a breeding program such as the present one, where genetic improvement is pursued for one target trait while constraining another essential trait within a desirable range. MT-LAS (Moeinizade et al. 2020) addresses a similar problem setting; however, it does not account for the progressive fixation of genotypes through repeated selfings, limiting its applicability to breeding programs for self-pollinated crops. CPS-MT utilizes the estimated marker effects and has been evaluated under several genetic relationships between two traits and their underlying causes, demonstrating its high effectiveness under various scenarios (Tables 1, 2, and Figure 5). Although this study focused on the analysis of two traits, a target and an essential trait, the CPS-MT framework can be directly applied to multi-trait constraints. A demonstration simulation with three traits (one target trait and two essential traits) is provided in Supplementary File S1, confirming that CPS-MT performs correctly under more than one essential trait constraint.

### Genetic variance and prediction accuracy

Genetic variance and diversity are sources of genetic improvements (Sanchez et al. 2023; Sanchez et al. 2024). Crosses selected based on UC tend to produce populations with greater genetic variance than those selected based on GEBV (Sakurai et al. 2024). The present results are consistent with these findings. CPS-MT maintained a higher genetic variance for the target trait (Trait1) compared to PCS across all seven scenarios (Figure 6). In addition, repeated selection has been reported to reduce genetic variance, leading to a decline in the prediction accuracy of GP models (Jannink 2010). In particular, in “No Relation” and “Nonlinear1” scenarios, CPS-MT maintained higher genetic variance than PCS did (Figure 6). These differences in genetic variance may contribute to the differences in prediction accuracy observed between CPS-MT and PCS (Figure 7). The maintenance of high genetic variance for the target trait under CPS-MT contributes to a higher prediction accuracy and continuous genetic improvement, which represents one of the strengths of CPS-MT (Figures 6 and 7).

Even in cases such as “Nonlinear1” and “Nonlinear2,” where nonlinear relationships exist between the genotypic values of Trait1 and Trait2, no substantial reduction in the prediction accuracy for Trait1 was observed compared with the “No Relation” and “Positive” scenarios under CPS-MT (Figure 7). Although the segregation of progeny no longer follows the bivariate Gaussian distribution described in Eq.24 under such nonlinear relationships, the superiority of CPS-MT over PCS was still confirmed (Tables 1, 2, and Figure 5). Therefore, CPS-MT can be effectively applied, regardless of the nature of the genetic relationships between traits.

### Prediction accuracy of genetic variance

In the early generations of this breeding program, the prediction accuracies of the genetic (co)variances in the Inbred8 population (*σ*_(11)_, *σ*_(22)_, and *σ*_(12)_) were low; however, they increased as the generations progressed (Figure 8). This improvement was due to an increase in the accuracy of the estimated marker effects (Figure S6). It has been reported that the prediction accuracy of genetic (co)variances in progeny populations depends on the accuracy of the estimated marker effects (Lehermeier et al. 2017; Oget-ebrad et al. 2024). In addition, achieving a high prediction accuracy in GP models requires a high degree of genetic similarity between the training and test populations (Lee et al. 2017a; Lee et al. 2017b; Jannink et al. 2023). In this breeding program, the high genetic similarity between the 100 individuals used to update the GP model in the Field Trial Component and the 150 individuals in the Population Improvement Component likely contributed to the high prediction accuracy (Figure 1). UC, which utilizes genetic variance in the progeny population, has been reported to become more effective as the accuracy of the estimated marker effects increases (des Déserts et al. 2023). Therefore, the improved accuracy of both the estimated marker effects and genetic (co)variances across generations is likely a factor contributing to the increasing superiority of CPS-MT over PCS (Figure 5).

### Usefulness and limitations of CPS-MT

In this study, we propose a new breeding strategy, CPS-MT, which aims to achieve genetic improvement of a target trait while maintaining the genotypic values of essential traits within desirable ranges. Various relationships exist between the target and essential traits, including positive correlations (Rincker et al. 2014; Assefa et al. 2018), negative correlations (Rosales-Serna et al. 2004), and nonlinear relationships (Ramírez-Oliveras et al. 1997; Tripathi et al. 2004; Craufurd and Wheeler 2009; Shah et al. 2017). These relationships are caused by factors such as pleiotropy and spurious pleiotropy (Reinert 2022; Dwivedi et al. 2024). In this study, seven scenarios reflecting these various relationships were designed, and the usefulness of CPS-MT was evaluated under each scenario. The results confirmed that CPS-MT effectively achieved genetic improvement of the target trait, while maintaining the genotypic value of the essential trait within the desirable range across all scenarios (Figure 5). These findings suggest that CPS-MT can be expected to serve as an effective breeding strategy for various situations. Despite these advantages, CPS-MT has several methodological limitations. This study assumed that the genotypic values in the progeny population produced from each cross followed a Gaussian distribution. When traits are controlled by a small number of QTLs and the distribution deviates from the Gaussian distribution, it becomes challenging to accurately estimate the potential of the parents. Moreover, gene-gene interactions (epistasis) were not considered in this breeding simulation, leaving the issue of more complex breeding programs unresolved. To further enhance the practicality of CPS-MT, flexible problem settings that meet the requirements of actual breeding programs and validation under more realistic conditions are required.

## Data availability

All datasets and source codes for the breeding simulations are available from the repository on GitHub, “https://github.com/Sakuraikengo/CPSMT.”

## Acknowledgments

The authors thank Dr. Akito Kaga for his advice for this study.

## Funding

This work was supported by JST SPRING Grant Number JPMJSP2108, JSPS KAKENHI Grant Number 25K23634, and JSPS International Leading Research Grant Number 22K21352.

## Conflicts of interest

The authors declare no conflict of interest.

## Literature cited

Addisu M, Snape JW, Simmonds JR, Gooding MJ. 2010. Effects of reduced height (Rht) and photoperiod insensitivity (Ppd) alleles on yield of wheat in contrasting production systems. Euphytica. 172(2):169–181. doi:10.1007/s10681-009-0025-2.

Allier A, Lehermeier C, Charcosset A, Moreau L, Teyssèdre S. 2019a. Improving short-and long-term genetic gain by accounting for within-family variance in optimal cross-selection. Front Genet. 10:1006. doi:10.3389/fgene.2019.01006.

Allier A, Moreau L, Charcosset A, Teyssèdre S, Lehermeier C. 2019b. Usefulness criterion and post-selection parental contributions in multi-parental crosses: Application to polygenic trait introgression. G3 Genes, Genomes, Genet. 9(5):1469–1479. doi:10.1534/g3.119.400129.

Allier A, Teyssèdre S, Lehermeier C, Moreau L, Charcosset A. 2020. Optimized breeding strategies to harness genetic resources with different performance levels. BMC Genomics. 21(1):349. doi:10.1186/s12864-020-6756-0.

Assefa Y, Bajjalieh N, Archontoulis S, Casteel S, Davidson D, Kovács P, Naeve S, Ciampitti IA. 2018. Spatial Characterization of Soybean Yield and Quality (Amino Acids, Oil, and Protein) for United States. Sci Rep. 8(1):1–11. doi:10.1038/s41598-018-32895-0.

Atanda SA, Bandillo N. 2024. Genomic-inferred cross-selection metrics for multi-trait improvement in a recurrent selection breeding program. Plant Methods. 20(1):133. doi:10.1186/s13007-024-01258-4.

Beckett TJ, Rocheford TR, Mohammadi M. 2019. Reimagining maize inbred potential: Identifying breeding crosses using genetic variance of simulated progeny. Crop Sci. 59(4):1457–1468. doi:10.2135/cropsci2018.08.0508.

Casebow R, Hadley C, Uppal R, Addisu M, Loddo S, Kowalski A, Griffiths S, Gooding M. 2016. Reduced height (Rht) alleles affect wheat grain quality. PLoS One. 11(5):1–20. doi:10.1371/journal.pone.0156056.

Chen B, Chai C, Duan M, Yang X, Cai Z, Jia J, Xia Q, Luo S, Yin L, Li Y, et al. 2024. Identification of quantitative trait loci for lodging and related agronomic traits in soybean (Glycine max [L.] Merr.). BMC Genomics. 25(1):1–16. doi:10.1186/s12864-024-10794-1.

Craufurd PQ, Wheeler TR. 2009. Climate change and the flowering time of annual crops. J Exp Bot. 60(9):2529–2539. doi:10.1093/jxb/erp196.

des Déserts AD, Durand N, Servin B, Goudemand-Dugué E, Alliot JM, Ruiz D, Charmet G, Elsen JM, Bouchet S. 2023. Comparison of genomic-enabled cross selection criteria for the improvement of inbred line breeding populations. G3 Genes, Genomes, Genet. 13(11):1–15. doi:10.1093/g3journal/jkad195.

Dwivedi SL, Heslop-Harrison P, Amas J, Ortiz R, Edwards D. 2024. Epistasis and pleiotropy-induced variation for plant breeding. Plant Biotechnol J. 22(10):2788–2807. doi:10.1111/pbi.14405.

Endelman JB. 2011. Ridge Regression and Other Kernels for Genomic Selection with R Package rrBLUP. Plant Genome. 4(3):250–255. doi:10.3835/plantgenome2011.08.0024.

Flower DJ. 1996. Physiological and morphological features determining the performance of the sorghum landraces of northern Nigeria. Exp Agric. 32(2):129–141. doi:10.1017/s0014479700026041.

Gaynor RC, Gorjanc G, Bentley AR, Ober ES, Howell P, Jackson R, Mackay IJ, Hickey JM. 2017. A two-part strategy for using genomic selection to develop inbred lines. Crop Sci. 57(5):2372–2386. doi:10.2135/cropsci2016.09.0742.

Hamazaki K, Iwata H. 2020. Rainbow: Haplotype-based genome-wide association study using a novel SNP-set method. PLoS Comput Biol. 16(2):1–17. doi:10.1371/journal.pcbi.1007663.

Hamazaki K, Iwata H. 2024. AI-assisted selection of mating pairs through simulation-based optimized progeny allocation strategies in plant breeding. Front Plant Sci. 15:1361894. doi:10.3389/fpls.2024.1361894.

Hazel LN. 1943. The Genetic Basis for Constructing Selection Indexes. Genetics. 28(6):476–490. doi:10.1093/genetics/28.6.476.

Hazel LN, Lush JL. 1942. The efficiency of three methods of selection. J Hered. 33(11):393–399. doi:10.1093/oxfordjournals.jhered.a105102.

Ibrahim AU. 2019. Genetic variability, Correlation and Path analysis for Yield and yield components in F6 generation of Wheat (Triticum aestivum Em. Thell.). IOSR J Agric Vet Sci. 12(1):17–23. doi:10.9790/2380-1201011723.

Jannink JL. 2010. Dynamics of long-term genomic selection. Genet Sel Evol. 42(1):1–11. doi:10.1186/1297-9686-42-35.

Jannink JL, Astudillo R, Frazier P. 2023. Insight into a two-part plant breeding scheme through Bayesian optimization of budget allocations. Crop Sci. 65(1):e21124. doi:10.1002/csc2.21124.

Kajiya-Kanegae H, Nagasaki H, Kaga A, Hirano K, Ogiso-Tanaka E, Matsuoka M, Ishimori M, Ishimoto M, Hashiguchi M, Tanaka H, et al. 2021. Whole-genome sequence diversity and association analysis of 198 soybean accessions in mini-core collections. DNA Res. 28(1):1–13. doi:10.1093/dnares/dsaa032.

Kinoshita S, Sakurai K, Hamazaki K, Tsusaka T, Sakurai M, Shirasawa K, Isobe S, Iwata H. 2025. Optimization of crossing strategy based on the usefulness criterion in interpopulation crosses considering different marker effects among populations. Theor Appl Genet. 138(7):1–16. doi:10.1007/s00122-025-04935-7.

Lee SH, Clark S, Van Der Werf JHJ. 2017. Estimation of genomic prediction accuracy from reference populations with varying degrees of relationship. PLoS One. 12(12):e0189775. doi:10.1371/journal.pone.0189775.

Lee SH, Weerasinghe WMSP, Wray NR, Goddard ME, Van Der Werf JHJ. 2017. Using information of relatives in genomic prediction to apply effective stratified medicine. Sci Rep. 7:1–13. doi:10.1038/srep42091.

Lehermeier C, de los Campos G, Wimmer V, Schön CC. 2017. Genomic variance estimates: With or without disequilibrium covariances? J Anim Breed Genet. 134(3):232–241. doi:10.1111/jbg.12268.

Lehermeier C, Teyssèdre S, Schön CC. 2017. Genetic gain increases by applying the usefulness criterion with improved variance prediction in selection of crosses. Genetics. 207(4):1651–1661. doi:10.1534/genetics.117.300403.

Li R, Li M, Ashraf U, Liu S, Zhang J. 2019. Exploring the relationships between yield and yield-related traits for rice varieties released in china from 1978 to 2017. Front Plant Sci. 10:543. doi:10.3389/fpls.2019.00543.

Meuwissen THE, Hayes BJ, Goddard ME. 2001. Prediction of total genetic value using genome-wide dense marker maps. Genetics. 157(4):1819–1829. doi:10.1093/genetics/157.4.1819.

Misztal I, Lourenco D. 2024. Potential negative effects of genomic selection. J Anim Sci. 102:skae155. doi:10.1093/jas/skae155.

Moeinizade S, Kusmec A, Hu G, Wang L, Schnable PS. 2020. Multi-trait genomic selection methods for crop improvement. Genetics. 215(4):931–945. doi:10.1534/genetics.120.303305.

Mohammadi M, Tiede T, Smith KP. 2015. Popvar: A genome-wide procedure for predicting genetic variance and correlated response in biparental breeding populations. Crop Sci. 55(5):2068–2077. doi:10.2135/cropsci2015.01.0030.

Navabi A, Iqbal M, Strenzke K, Spaner D. 2006. The relationship between lodging and plant height in a diverse wheat population. Can J Plant Sci. 86(3):723–726. doi:10.4141/P05-144.

Neyhart JL, Smith KP. 2019. Validating genomewide predictions of genetic variance in a contemporary breeding program. Crop Sci. 59(3):1062–1072. doi:10.2135/cropsci2018.11.0716.

Oget-ebrad C, Heumez E, Duchalais L, Goudemand-dugué E, Oury F. 2024. Validation of cross progeny variance genomic prediction using simulations and experimental data in winter elite bread wheat. Theor Appl Genet. 137(10):226. doi:10.1007/s00122-024-04718-6.

Ramírez-Oliveras G, Stutte CA, Orengo-Santiago E. 1997. Hydrogen ion efflux differences in soybean roots associated with yields. J Agric Univ Puerto Rico. 81(3–4):159–180. doi:10.46429/jaupr.v81i3-4.3640.

Reinert S. 2022. Quantitative genetics of pleiotropy and its potential for plant sciences. J Plant Physiol. 276:153784. doi:10.1016/j.jplph.2022.153784.

Rincker K, Nelson R, Specht J, Sleper D, Cary T, Cianzio SR, Casteel S, Conley S, Chen P, Davis V, et al. 2014. Genetic Improvement of U.S. Soybean in Maturity Groups II, III, and IV. Crop Sci. 54(4):1419–1432. doi:10.2135/cropsci2013.10.0665.

Rosales-Serna R, Kohashi-Shibata J, Acosta-Gallegos JA, Trejo-López C, Ortiz-Cereceres J, Kelly JD. 2004. Biomass distribution, maturity acceleration and yield in drought-stressed common bean cultivars. F Crop Res. 85(2–3):203–211. doi:10.1016/S0378-4290(03)00161-8.

Sakurai K, Hamazaki K, Inamori M, Kaga A, Iwata H. 2024. Cross potential selection: a proposal for optimizing crossing combinations in recurrent selection using the usefulness criterion of future inbred lines. G3 Genes, Genomes, Genet. 14(11):jkae224. doi:10.1093/g3journal/jkae224.

Sanchez D, Allier A, Ben Sadoun S, Mary-Huard T, Bauland C, Palaffre C, Lagardère B, Madur D, Combes V, Melkior S, et al. 2024. Assessing the potential of genetic resource introduction into elite germplasm: a collaborative multiparental population for flint maize. Theor Appl Genet. 137(1):19. doi:10.1007/s00122-023-04509-5.

Sanchez D, Sadoun S, Mary-Huard T, Allier A, Moreau L, Charcosset A. 2023. Improving the use of plant genetic resources to sustain breeding programs’ efficiency. Proc Natl Acad Sci. 120(14):e2205780119. doi:10.1073/pnas.2205780119.

Schnell FW, Utz HF. 1975. F1-Leistung und Elternwahl in der Züchtung von Selbstbefruchtern. In: Bericht über die Arbeitstagung der Vereinigung österreichischer Pflanzenzüchter. Gumpenstein. p. 243–248.

Shah AN, Tanveer M, Rehman A ur, Anjum SA, Iqbal J, Ahmad R. 2017. Lodging stress in cereal— effects and management: an overview. Environ Sci Pollut Res. 24(6):5222–5237. doi:10.1007/s11356-016-8237-1.

Strandén I, Garrick DJ. 2009. Derivation of equivalent computing algorithms for genomic predictions and reliabilities of animal merit. J Dairy Sci. 92(6):2971–2975. doi:10.3168/jds.2008-1929.

Tiede T, Kumar L, Mohammadi M, Smith KP. 2015. Predicting genetic variance in bi-parental breeding populations is more accurate when explicitly modeling the segregation of informative genomewide markers. Mol Breed. 35(10):1–13. doi:10.1007/s11032-015-0390-6.

Tripathi SC, Sayre KD, Kaul JN, Narang RS. 2004. Lodging behavior and yield potential of spring wheat (Triticum aestivum L.): Effects of ethephon and genotypes. F Crop Res. 87(2–3):207–220. doi:10.1016/j.fcr.2003.11.003.

VanRaden PM. 2008. Efficient methods to compute genomic predictions. J Dairy Sci. 91(11):4414–4423. doi:10.3168/jds.2007-0980.

Wartha CA, Lorenz AJ. 2024. Genomic predictions of genetic variances and correlations among traits for breeding crosses in soybean. Heredity (Edinb). 133(3):173–185. doi:10.1038/s41437-024-00703-3.

Wimmer V, Lehermeier C, Albrecht T, Auinger HJ, Wang Y, Schön CC. 2013. Genome-wide prediction of traits with different genetic architecture through efficient variable selection. Genetics. 195(2):573–587. doi:10.1534/genetics.113.150078.

Wu DH, Chen CT, Yang M Der, Wu YC, Lin CY, Lai MH, Yang CY. 2022. Controlling the lodging risk of rice based on a plant height dynamic model. Bot Stud. 63(1). doi:10.1186/s40529-022-00356-7.

Yamada T, Hajika M, Yamada N, Hirata K, Okabe A, Oki N, Takahashi K, Seki K, Okano K, Fujita Y, et al. 2012. Effects on flowering and seed yield of dominant alleles at maturity loci E2 and E3 in a Japanese cultivar, Enrei. Breed Sci. 61(5):653–660. doi:10.1270/jsbbs.61.653.

Yao J, Zhao D, Chen X, Zhang Y, Wang J. 2018. Use of genomic selection and breeding simulation in cross prediction for improvement of yield and quality in wheat (Triticum aestivum L.). Crop J. 6(4):353–365. doi:10.1016/j.cj.2018.05.003.

Zhou J, Beche E, Vieira CC, Yungbluth D, Zhou J, Scaboo A, Chen P. 2022. Improve Soybean Variety Selection Accuracy Using UAV-Based High-Throughput Phenotyping Technology. Front Plant Sci. 12:768742. doi:10.3389/fpls.2021.768742.

Zhu G, Li G, Wang D, Yuan S, Wang F. 2016. Changes in the lodging-related traits along with rice genetic improvement in China. PLoS One. 11(7):1–14. doi:10.1371/journal.pone.0160104.

